# A complete collagen IV fluorophore knock-in toolkit reveals α-chain diversity in basement membrane

**DOI:** 10.1101/2024.12.14.628508

**Authors:** Sandhya Srinivasan, Willam Ramos-Lewis, Mychel R.P.T. Morais, Qiuyi Chi, Adam W. J. Soh, Emily Williams, Rachel Lennon, David R. Sherwood

**Affiliations:** Department of Biology, Duke University, 130 Science Drive, Box 90338, Durham, NC 27708, USA; Wellcome Centre for Cell-Matrix Research, Division of Cell-Matrix Biology and Regenerative Medicine, School of Biological Sciences, Faculty of Biology Medicine and Health, The University of Manchester, Manchester Academic Health Science Centre, Manchester, M13 9PT, UK

## Abstract

The type IV collagen triple helix, composed of three ⍺-chains, is a core basement membrane (BM) component that assembles into a network within BMs. Endogenous tagging of all ⍺-chains with genetically encoded fluorophores has remained elusive, limiting our understanding of this crucial BM component. Through genome editing, we show that the C-termini of the *C. elegans* type IV collagen ⍺-chains EMB-9 and LET-2 can be fused to a variety of fluorophores to create a strain toolkit with wild-type health. Using quantitative imaging, our results suggest a preference for LET-2-LET-2-EMB-9 trimer construction, but also tissue-specific flexibility in trimers assembled driven by differences in ⍺-chain expression levels. By tagging *emb-9* and *let-2* mutants that model human Gould Syndrome, a complex multi-tissue disorder, we further discover defects in extracellular accumulation and turnover that might help explain disease pathology. Together, our findings identify a permissive tagging site that will allow diverse studies on type IV collagen regulation and function in animals.

**Summary:** Srinivasan et al., construct a collagen IV fluorophore knock-in toolkit in *C. elegans* using a newly identified permissive genome editing site and reveal tissue-specific α-chain diversity and basement membrane turnover defects in collagen IV mutants modeling human COL4A1/A2 (Gould) syndrome.

## Introduction

Basement membranes (BMs) are dense, sheet-like extracellular matrices that surround and support most tissues (Fidler et al., 2018; Jayadev and Sherwood, 2017a). Type IV collagen is a triple helical protomer and a core BM component that forms a self-associating network in BMs. The type IV collagen BM network confers resistance to mechanical loads placed on tissues (Fidler et al., 2018; Keeley et al., 2020; Morrissey and Sherwood, 2015) and type IV collagen genetic loss leads to embryonic lethality at the onset of muscle contraction in mice, *Drosophila*, and *C. elegans* (Borchiellini et al., 1996; Gupta et al., 1997; Poschl et al., 2004). Precise deposition and remodeling of type IV collagen is also required to properly shape tissues, such as the *Drosophila* egg chamber, central nervous system, and wing disc and the mouse salivary gland (Crest et al., 2017; Harunaga et al., 2014; Isabella and Horne-Badovinac, 2016; Morrissey and Sherwood, 2015; Pastor-Pareja and Xu, 2011). Additionally, type IV collagen networks serve as scaffolds that bind signaling ligands and other BM components that direct cell differentiation and guide cell migration (Brown et al., 2017; Morrissey and Sherwood, 2015; Parkin et al., 2011; Sherwood, 2021; Wang et al., 2008). Further emphasizing its importance to tissue function, pathogenic variants in type IV collagen are linked to at least ten human genetic disorders that affect muscle, kidney, brain and vasculature (Fidler et al., 2018). Additionally, increased type IV collagen BM accumulation is associated with the decline of vascular function in diabetes, Alzheimer’s, and during aging. (Baum and Bigler, 2016; Candiello et al., 2010; Fidler et al., 2018; Howe et al., 2020; Karttunen et al., 1986; Marshall, 2016; Thomsen et al., 2017; Uspenskaia et al., 2004).

The triple helical type IV collagen protomer is composed of three ⍺-chains, which have a non-collagenous 7S domain at the N-terminus, a central interrupted Gly-X-Y repeat region (X and Y can be any amino acid, but is most often proline), and a non-collagenous NC1 domain at the C-terminus (Hudson et al., 1993; Hudson et al., 2003). Reconstitution studies have shown that the NC1 domain mediates α-chain selection for type IV collagen trimer assembly, which occurs in the endoplasmic reticulum (ER) (Boutaud et al., 2000; Dolz et al., 1988; Fidler et al., 2018). In-vitro studies have also revealed that lysyl- and prolyl-hydroxylases, along with the cofactor vitamin C, are needed for proper folding and trimerization (Kivirikko and Pihlajaniemi, 1998; Lamande and Bateman, 1999). Depleting prolyl-4-hydroxylase in *Drosophila* larvae results in secretion of type IV collagen in its monomeric form and an absence of BM localization, suggesting BM incorporation requires trimerization (Pastor-Pareja and Xu, 2011). Upon secretion into the extracellular space, two type IV collagen protomers connect through the C-terminal NC1 domains and four protomers connect through the N-terminal 7S domains (Anazco et al., 2016; Cummings et al., 2016). The interfaces of the NC1 and 7S connections are reinforced with covalent crosslinks (Cummings et al., 2016; Vanacore et al., 2009). Together, the NC1 dimer and 7S tetramer connections establish a polygonal patterned network of type IV collagen in BM. Electron microscopy studies have also revealed higher order lateral twisting interactions between neighboring protomers, which is thought to provide further stability to type IV collagen networks (Barnard et al., 1992; Ruben and Yurchenco, 1994; Yurchenco and Ruben, 1987).

Based on sequence similarity, type IV collagen α-chains have been divided into α1-like and α2-like groups. Most vertebrates have six α-chain encoding genes (*COL4A1-COL4A6* genes encoding the α1-α6 proteins). The α1-like chain group contains *COL4A1*, *COL4A3,* and *COL4A*5 genes and the α2-like group includes *COL4A*2, *COL4A*4, and *COL4A*6 (Khoshnoodi et al., 2008). *Caenorhabditis elegans* and *Drosophila melanogaster* each possess two type IV collagen α-chains, encoded by *emb-9* and *let-2* in *C. elegans* and *vkg* and *Cg25C* in *Drosophila* (Guo and Kramer, 1989; Natzle et al., 1982; Sibley et al., 1993; Yasothornsrikul et al., 1997). Sequence comparisons with mammalian NC1 domains have suggested that *emb-9* and *vkg* are more similar to ⍺1-like chains and *let-2* and *Cg25C* are more similar to ⍺2-like chains (Guo and Kramer, 1989; Summers et al., 2023).

Chromatography-based protein isolation studies in vertebrates have revealed approximately twice as much α1 compared to the α2 chain in several tissues (Mayne and Zettergren, 1980; Siebold et al., 1988; Trueb et al., 1982), which is consistent with the existence of the α1α1α2 heterotrimer. Characterization of type IV collagen hexamers (the NC1 domains of two cross-linked protomers) isolated from in vivo tissues have also supported the existence of α1α1α2, as well as α3α4α5 and α5α5α6 type IV collagen trimers (Borza et al., 2001; Boutaud et al., 2000; Khoshnoodi et al., 2008). These and other biochemical studies have led to the proposition that only three (out of a possible 56) specific combinations of vertebrate type IV collagen are assembled in vivo —α1α1α2, α3α4α5, and α5α5α6—with α1α1α2 being the most abundant and ubiquitous (Borza et al., 2001; Boutaud et al., 2000; Hudson et al., 1994; Hudson et al., 2003; Khoshnoodi et al., 2008).

While there is strong support for the preferential formation of α1α1α2, α3α4α5, and α5α5α6 type IV collagen trimers in mammalian tissues, there is also evidence that other combinations could exist. For example, isolation and characterization of type IV collagen hexamers, have not ruled out the existence of α1 homotrimers or α1α2α2 heterotrimers (Boutaud et al., 2000; Johansson et al., 1992; Siebold et al., 1988). Indeed, reconstitution experiments using bovine and recombinant NC1 α1-α5 monomers have found that α1 NC1 domains can form a homohexamer and that the α2 NC1 can assemble in a promiscuous manner with other NC1 domains (Boutaud et al., 2000). Further, rat carcinoma cells generate and secrete α1 homotrimers (Haralson et al., 1985). No studies have yet addressed the overall α-chain levels and possible trimer combinations found in *C. elegans.* Mutations in the *emb-9* or *let-2,* however, result in intracellular retention of the other (non-mutant) α-chain (Gupta et al., 1997), suggesting a preference for heterotrimer assembly prior to secretion. Recombinant expression of forced pre-determined NC1 trimers in *Drosophila* larvae has shown that only the Cg25C-Cg25C-Vkg NC1 heterotrimer combination is recruited to the BM. Yet, NC1 domains expressed as individual monomers in mammalian cells assemble in almost all possible combinations. Additionally, analysis of endogenous hexamers did not rule out formation of homotrimers (Summers et al., 2023). Together these observations suggest that flexibility in the α-chain composition of type IV collagen trimers might exist in BMs in vivo.

Endogenous tagging of BM components with genetically encoded fluorophores allows quantitative comparisons of component abundance at the tissue, cell, and subcellular level (Jayadev et al., 2022; Keeley et al., 2020). A key challenge for tagging type IV collagen is identifying a fluorophore insertion site common to all α-chains that does not disrupt type IV collagen assembly and function. To date, endogenous tagging in mice, *Drosophila* and *C. elegans* has predominantly used the *Drosophila* Vkg-GFP protein trap insertion site, which inserts GFP within the start of the 7S domain at the N-terminus (Morin et al., 2001). While insertion of fluorophores at this site is viable for Drosophila *vkg*, *C. elegans emb-9*, and mouse *COL4A2* (Keeley et al., 2020; Wuergezhen et al., 2025), insertion at this site is not viable in *C. elegans let-2* or mouse *COL4A1* (Jones et al., 2024; Keeley et al., 2020). A site within an interruption of the Gly-X–Y repeats in the collagenous domain of *C. elegans emb-9* has also been reported to be viable (Matsuo et al., 2019), but the health of this strain has not been fully explored.

Here, we use genome editing, fitness assays, and live cell imaging to compare sites for fluorophore insertion into the *C. elegans emb-9* and *let-2* type IV collagen α-chains. We show that fluorophore insertion at the Vkg-GFP site (renamed Internal Site 1, IS1) and an internal site within the collagenous domain of EMB-9 (IS2) lead to accumulation of the tagged α-chains in the ER and reduced organism fitness. In contrast, we use multiple fluorophores and show that insertion at the C-terminus of EMB-9 and LET-2 proteins leads to dramatically less intracellular retention and phenotypically wild-type fitness. Using these new strains with mass spectrometry on whole animals, we reveal there is approximately two-fold more LET-2 protein than EMB-9, suggesting predominantly a LET-2-LET-2-EMB-9 (LLE) heterotrimer. However, using quantitative live imaging of mNG tagged strains we find that the LET-2/EMB-9 protein ratio present at various BMs is not precisely 2:1, but instead is ∼1.7 on most tissues and at the spermathecal BM is over 2.2. These results suggest a preference for the LLE heterotrimer, but that there is also flexibility in α-chain trimer make-up. Using tissue specific translational reporters, RNAi knockdown, and overexpression, we provide evidence suggesting that α-chain availability during assembly can drive trimer diversity. Finally, we show that the C-terminal site for type IV collagen allows endogenous tagging of type IV collagen variants found in COL4A1/A2 syndrome (also known as Gould Syndrome). Through localization and photoconversion analysis, we find that in addition to defects in type IV collagen secretion, these mutant forms affect extracellular accumulation and turnover. Together, our findings identify a permissive site for tagging type IV collagen α-chains, which will allow wide ranging studies.

## Results

### Healthy C-terminal fluorophore knock-ins of EMB-9 and LET-2 type IV collagen ⍺-chains

The inability to comprehensively label and visualize type IV collagen α-chains in animals has limited our understanding of type IV collagen regulation, function, and composition in BM. Fluorophore insertion at the C-terminus through genome editing has successfully labelled 37 of the 56 BM components tagged in C. *elegans* to date, but has not yet been attempted for the type IV collagen α-chain encoding genes *emb-9* and *let-2* (Fan et al., 2020; Hastie et al., 2019; Jayadev et al., 2022; Keeley et al., 2020; Naegeli et al., 2017). Thus, we knocked-in genetically encoded fluorescent proteins with a 18 amino acid linker within the endogenous *emb-9* and *let-2* genes at the C-terminus (C site) of the protein encoding regions using CRISPR/Cas9 gene editing (Fig. 1 and Fig. S1 A). The resulting gene edited animals were homozygous viable and fertile. To compare the health of these new strains to animals with fluorescent proteins inserted at the IS1 (*Drosophila* Vkg*-*GFP site) and IS2 sites of type IV collagen (Fig. 1), we plated four L4 larvae from each strain onto a predetermined amount of food and recorded how long it took for the food to be depleted (time to starvation). This served as a collective estimate of growth, feeding, and fertility. Eight strains with six distinct knock-ins at the C site of *emb-9* and *let-2* were analyzed: *emb-9* and *let-2* each tagged with mNeonGreen (mNG) or mRuby2, strains harboring both *let-2* and *emb-9* tagged with mNG and mRuby2 and vice versa, *emb-9* tagged with photoconvertible mEos2 and *let-2* tagged with photoconvertible mMaple (Heppert et al., 2016; McEvoy et al., 2012; Zhang et al., 2012) (Fig. 1 and Fig. S1, A and B). All strains were indistinguishable from wild-type animals, apart from *let-2* tagged with mMaple, which showed a slight delay in the time to starvation (Table 1). In contrast, *emb-9* strains tagged at the IS1 site (mRuby2, mScarlet-I, mEos2, mNG) and IS2 site (mCherry) (Bindels et al., 2017; Heppert et al., 2016) all showed delays to starvation (Table 1, Fig. 1 and Fig. S1, A and B). It is notable that all fluorophore insertions at the IS1 site (Vkg site), which has been used most extensively to label type IV collagen (Keeley et al., 2020; Morin et al., 2001; Wuergezhen et al., 2025), caused delayed growth.

**Figure 1.**
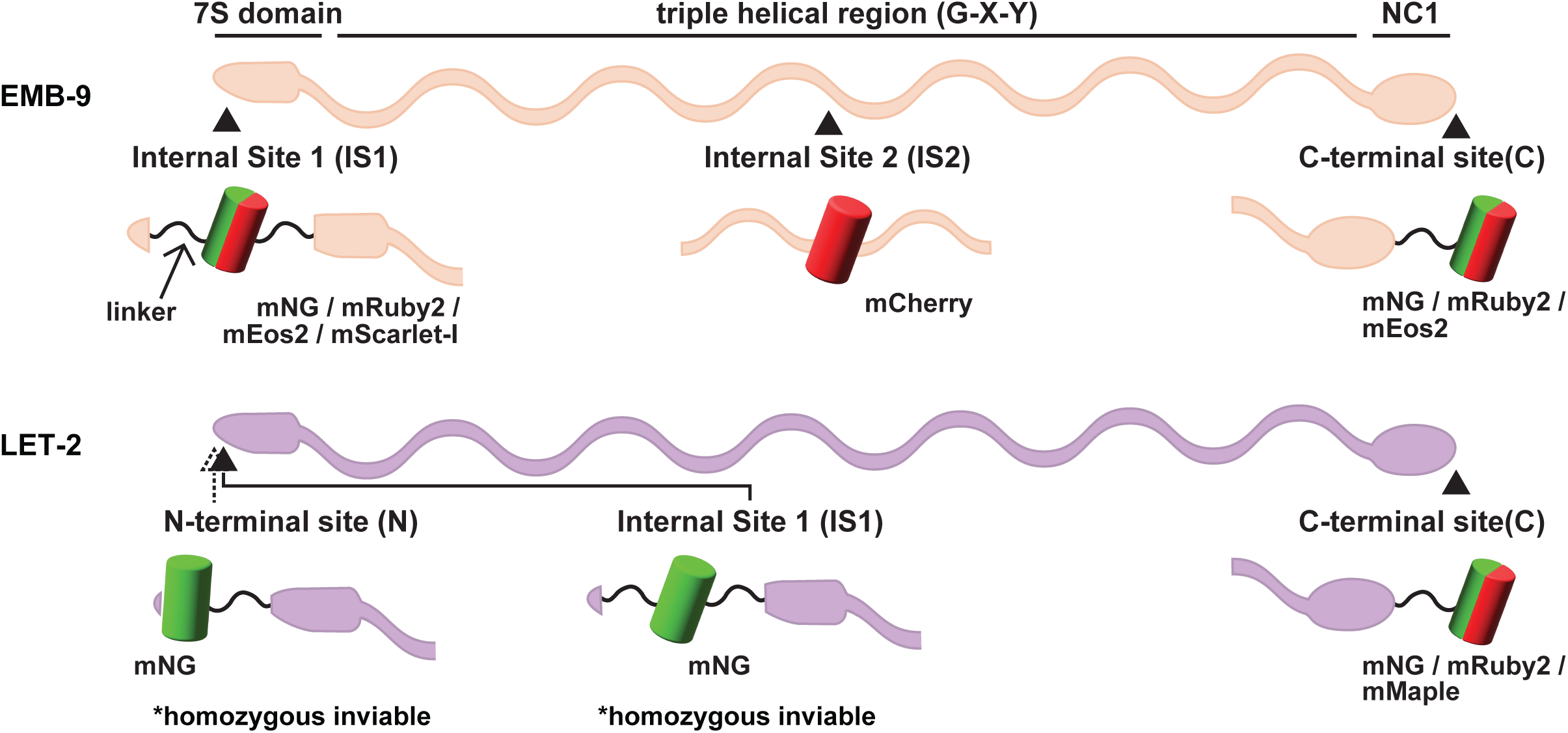
Fluorophore insertion sites in the *C*. *elegans* type IV collagen α-chains. The schematic illustrates *C*. *elegans* type IV collagen α-chains EMB-9 and LET-2 with the N-terminal 7S domains, the triple helical regions, and the C-terminal NC1 domains annotated above the proteins. Arrowheads mark fluorophore insertion sites. Each site is labelled with the fluorophores that were inserted at the location and whether the fluorophore was flanked by a linker sequence (thin black wavy lines, see Fig. S1 A). Insertions that were homozygous inviable are denoted.

**Table 1.**
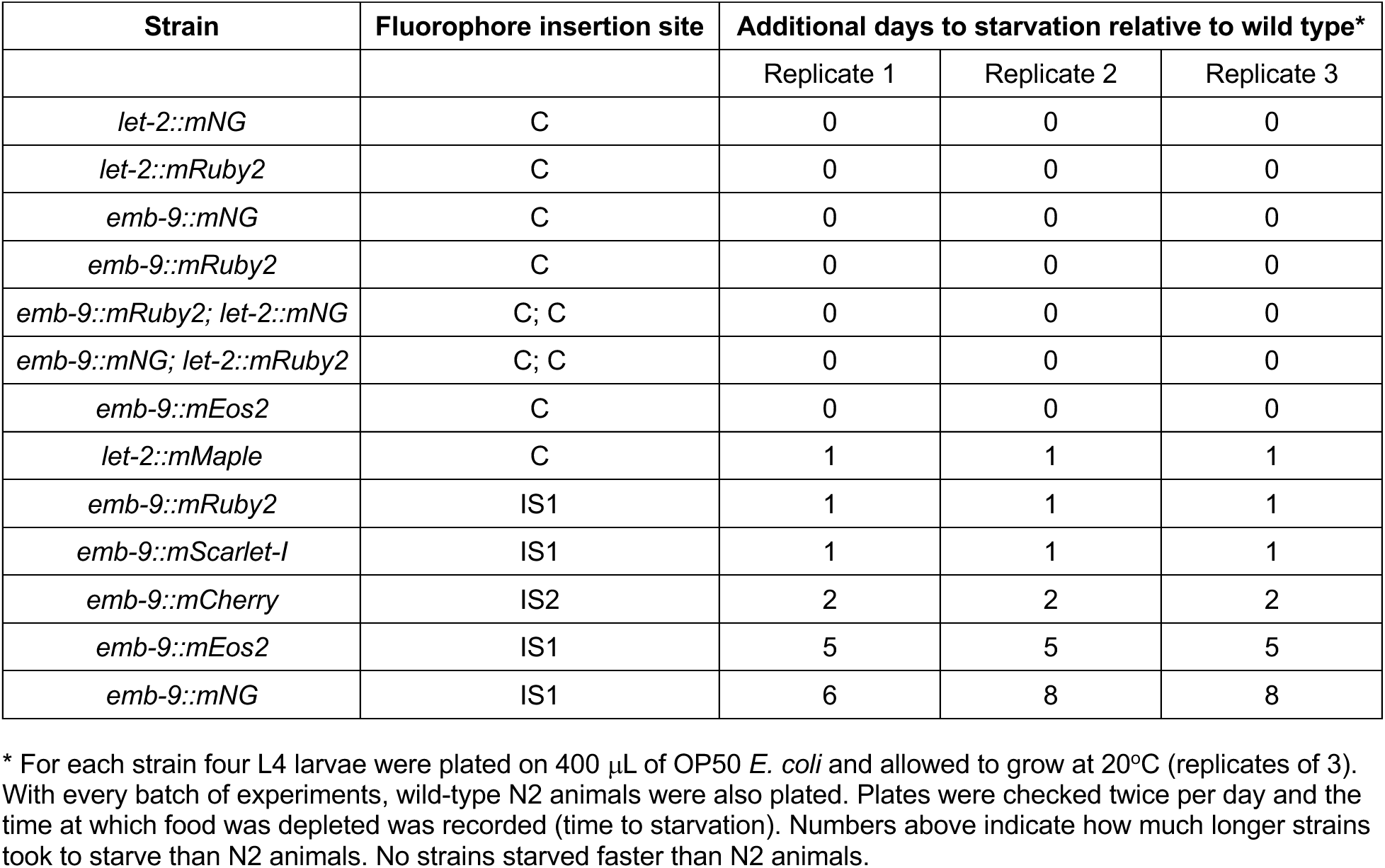
Health of fluorescently tagged type IV collagen strains.

We next wanted to compare protein trafficking and localization of the type IV collagen tagged in distinct sites. Type IV collagen is expressed in several tissues in *C. elegans*, but the predominant site of production is the *C. elegans* body wall muscle, where type IV collagen is made, secreted into BM and also trafficked from to many other tissue BMs (Graham et al., 1997; Morrissey et al., 2016). Confocal imaging of L1 staged larvae (0-12 h post hatch) showed large and extensive body wall muscle intracellular aggregates of EMB-9 tagged at the IS1 site with mNG, mEos2, mRuby2 or mScarlet-I, and aggregates of EMB-9 tagged at the IS2 site with mCherry (Fig. 2 A). In contrast, EMB-9 and LET-2 tagged with mNG or mRuby2 at the C site possessed few if any aggregates (Fig. 2 A). Previous immunolocalization studies also found limited intracellular aggregates of EMB-9 and LET-2 in wild-type L1 animals (Graham et al., 1997; Kang and Kramer, 2000), suggesting tagging at the C site permits normal intracellular trafficking. Imaging of worms tagged at the C site from the L1 through L4 stage and into the gravid adult revealed the gradual appearance of small type IV collagen aggregates starting at the L3 larval stage that continued to the adult (Fig. S2 A). Given the wild-type health of these strains, this might represent natural intracellular accumulation.

**Figure 2.**
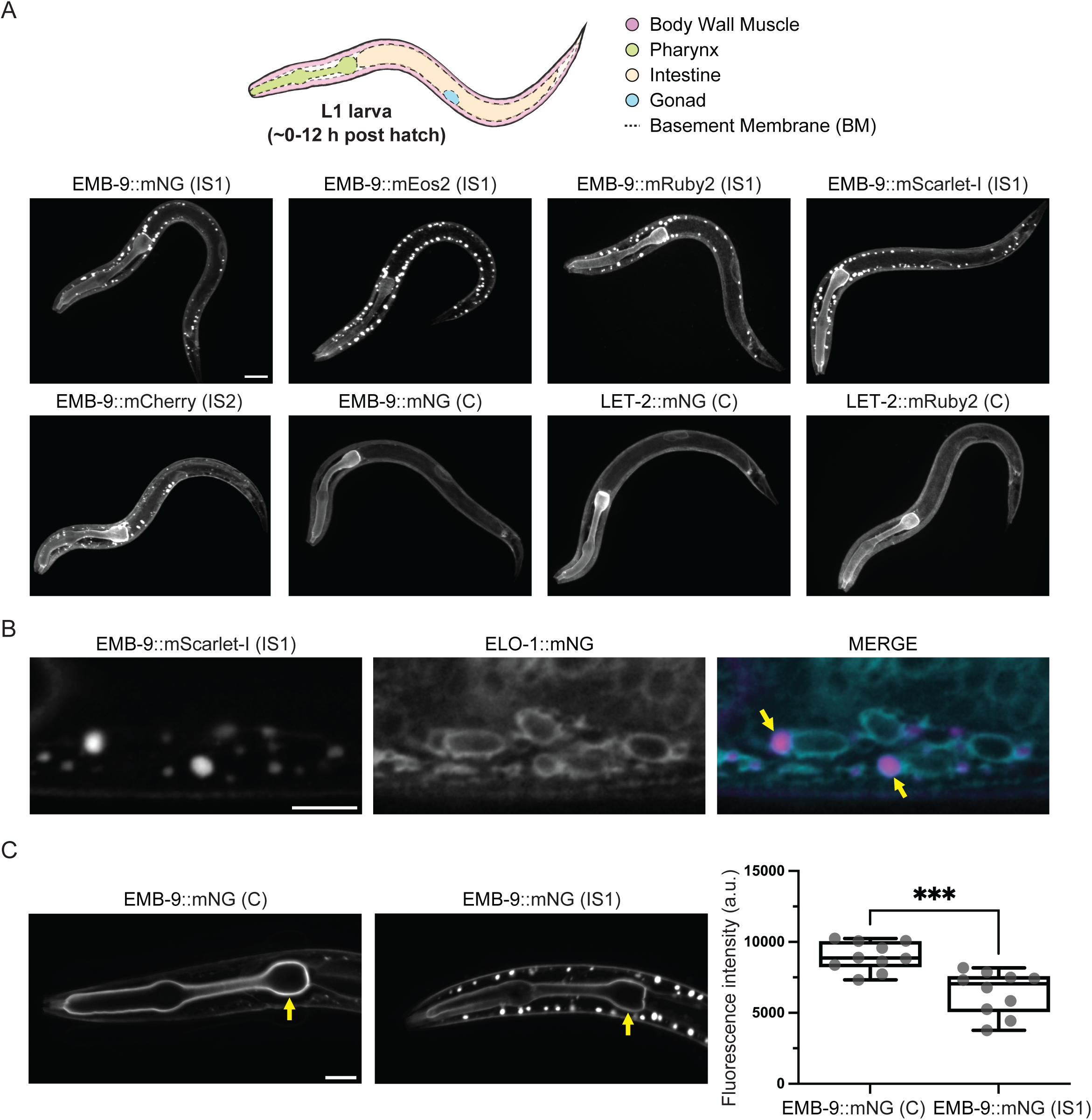
C-terminal EMB-9 and LET-2 fluorophore knock-ins are secreted and incorporated into BMs more efficiently than internal knock-ins. **(A)** Top: The schematic indicates location of tissues and surrounding BMs (dashed lines) analyzed in an L1 larva. Bottom: Representative max projections from confocal fluorescence z-stacks of L1 larvae with fluorescently tagged type IV collagen α-chains. Internally tagged α-chains aggregate in body wall muscles while C-terminally tagged α-chains do not (n = 10 animals per strain). The scale bar is 10 μm. **(B)** Single z-slice confocal images show that aggregates of EMB-9::mScarlet-I (IS1) (magenta in merge) are localized within ER membrane (visualized with ELO-1::mNG, cyan in merge) in L1 larvae. Yellow arrows indicate aggregates in the ER (100/100 aggregates localized to the ER, 10 aggregates counted per animal). The scale bar is 5 μm. **(C)** Left: Single z-slice confocal images show reduced EMB-9::mNG (IS1) localized to the pharyngeal BM compared to EMB-9::mNG (C) in L1 larvae (∼ 28% reduction, yellow arrows indicate pharyngeal BM). Right: Boxplots show mean fluorescence intensity measurements (a.u.) of the pharyngeal BM surrounding the proximal bulb (median of data denoted by horizontal line within each box for this and all subsequent boxplots) for EMB-9::mNG (C) versus EMB-9::mNG (IS1) animals (n = 10 animals each, *** P < 0.001, unpaired two-tailed Student’s t-test). The scale bar is 10 μm.

Type IV collagen folding and trimerization occurs in the ER, which has quality control mechanisms that retain misfolded proteins (Ellgaard and Helenius, 2003). We thus hypothesized that the type IV collagen aggregates found in strains tagged at the IS1 and IS2 sites were likely localized to the ER. Colocalization of type IV collagen aggregates tagged at the IS1 site (EMB-9::mScarlet-I (IS1)) with an ER marker (ELO-1::mNG (Park et al., 2024)), revealed that the EMB-9::mScarlet-I (IS1) aggregates were always enclosed by ER membrane (Fig. 2 B). Similarly, EMB-9::mCherry (IS2) aggregates were also contained within the ER (Fig. S2 B). This suggests that fluorophore insertion at the IS1 or IS2 site interferes with proper protein folding or trimerization, leading to ER accumulation and possibly reduced secretion. Consistent with this idea, measurements at the pharyngeal BM in L1 larvae showed that less EMB-9::mNG (IS1) localized to the BM compared to EMB-9::mNG (C) (Fig. 2 C). Decreased EMB-9::mNG (IS1) in the BM, might also be caused by alterations in BM on-off dynamics, often referred to as turnover (Stramer and Sherwood, 2024). However, fluorescence recovery in a photobleached region of EMB-9::mNG (IS1) compared with EMB-9::mNG (C) and LET-2::mNG (C), revealed similar recovery dynamics: LET-2::mNG (C) recovered ∼32% and both EMB-9::mNG (C) and EMB-9::mNG (IS1) recovered ∼25% of bleached fluorescence over a 5 h period (Fig. S2 C). Taken together, the wild-type health and greatly reduced intracellular aggregation, offers compelling evidence that the C-terminus is a permissive site for fluorescently tagging type IV collagen α-chains.

### LET-2 is approximately twice as abundant as EMB-9 in whole animals

We next wanted to determine the overall levels of EMB-9 and LET-2 proteins in the extracellular matrix of whole animals using the C site tagged strains as material for liquid chromatography (LC) coupled with tandem mass spectrometry (MS/MS). Comparisons of abundance of distinct proteins using LC-MS/MS is usually challenging because differences in the physical and chemical properties of distinct peptides can affect processing and detection (Bantscheff et al., 2007; Xie et al., 2011). However, the tagging of LET-2 and EMB-9 with the same fluorophore allowed us to use the fluorophore proteins as a proxy of LET-2 and EMB-9 abundance to determine their levels using label-free quantitative proteomics. We collected mixed populations of L1-L3 larval staged worms of animals expressing the knock-in proteins EMB-9::mRuby2 (C); LET-2::mNG (C) (strain 1) and EMB-9-mNG (C); LET-2::mRuby2 (C) (strain 2) and chemically fractionated them to enrich for extracellular matrix proteins as described previously (Morais et al., 2022; Morais et al., 2023). Measuring the protein abundance of mRuby2 in the extracellular matrix enriched fraction of these strains using LC-MS/MS revealed an ∼2-fold higher level of LET-2 compared to EMB-9 (Fig. 3 A). Although mNG was not detected in the matrix fraction, we found that in the cellular fraction the levels of mNG indicated LET-2 was ∼1.7 fold higher than EMB-9::mNG (Fig. S3 A). Similar results were observed in the cellular fraction for LET-2 and EMB-9 tagged with mRuby2 (Fig. S3 B). Finally, comparison of EMB-9 to EMB-9 and LET-2 to LET-2 protein levels between strain 1 [EMB-9::mRuby2 (C); LET-2::mNG (C)] and strain 2 [EMB-9-mNG (C); LET-2::mRuby2 (C)] matrix fractions confirmed that the protein abundances between the two strains were comparable (Fig. S3, C and D). Taken together, these studies reveal that LET-2 is present at approximately twice the levels of EMB-9 in larval stage *C. elegans*.

**Figure 3.**
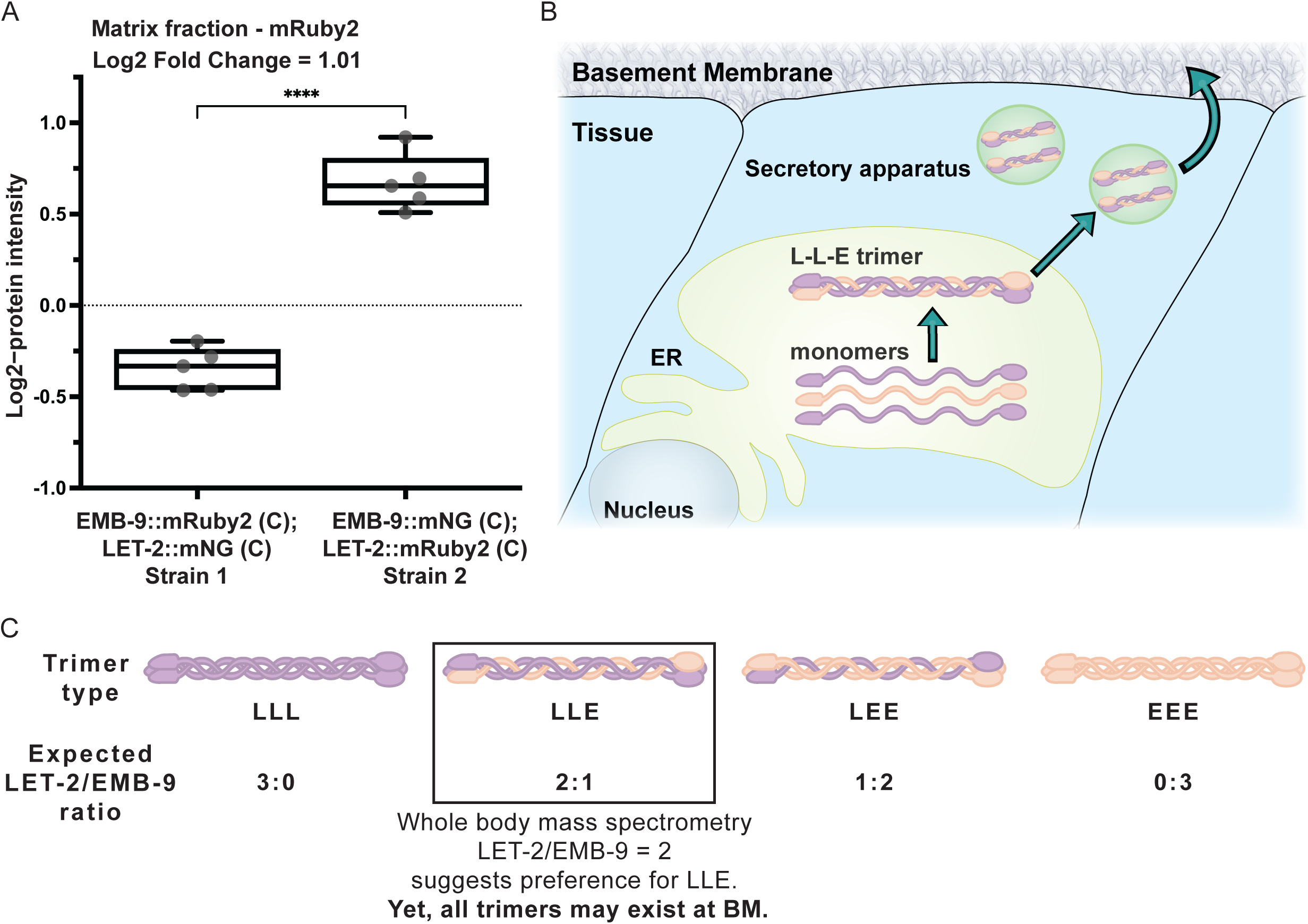
LET-2 is the more abundant type IV collagen α-chain. **(A)** Boxplots display the mRuby2 abundance represented by the log_2_-protein intensity (a.u.) determined using LC-MS/MS. The samples displayed are the larval matrix fractions of Strain 1 [EMB-9::mRuby2 (C); LET-2::mNG (C)] and Strain 2 [EMB-9-mNG (C); LET-2::mRuby2 (C)]. The log_2_-fold change from the comparison between strains is listed above the graph. mRuby2 (LET-2) was ∼2-times more abundant in Strain 2 than mRuby2 (EMB-9) in Strain 1 (n = 5 samples for each strain, P < 0.0001, unpaired two-tailed Student’s t-test). **(B)** The schematic demonstrates trimerization of *C. elegans* EMB-9 and LET-2 α-chains in the ER prior to secretion into the BM. **(C)** A schematic showing possible types of Collagen IV trimers and the LET-2/EMB-9 ratios expected for each timer type. Mass spectrometry on whole animals suggests a preference for the LLE trimer type but does not rule out different trimers being made and localized to BM.

### LET-2/EMB-9 α-chain BM ratios are not precisely 2:1 and vary between tissues

The 2:1 whole body protein ratio of LET-2/EMB-9, suggested a LET-2-LET-2-EMB-9 (LLE) heterotrimer combination is assembled in the ER and is present in BMs (Fig. 3 B and C). We thus next wanted to use our endogenously tagged EMB-9 and LET-2 strains to determine if a 2:1 LET-2/EMB-9 ratio is indeed present in BMs. We measured the average fluorescence intensity of LET-2::mNG (C) and EMB-9::mNG (C) in a specific region of BM in the pharynx (feeding organ), the body wall muscle, the distal tip cell (DTC, the cell that leads gonad migration), and the spermatheca (where oocyte fertilization and ovulation occurs) (Fig. 4 A) (Gieseler et al., 2017; Keeley et al., 2020; Kelley and Cram, 2019; Sherwood and Plastino, 2018). The ratio of LET-2/EMB-9 across the BMs of these tissues in L4.8 - L4.9 worms (1 h window in larval development) (Mok et al., 2015) revealed an average ratio of ∼1.7 within the pharynx, muscle and DTC, but a significantly greater ratio of ∼2.2 in the spermathecal BM (Fig. 4 A and B; Fig. S4 A). While all tissue BMs had LET-2/EMB-9 ratio close to 2:1, strongly suggesting the presence and perhaps preference for the LLE trimer, ratios of LET-2/EMB-9 lower than 2:1 indicated the existence of LEE or EEE or both trimers. Further, the ratio of greater than 2:1 in the spermathecal BM implied the presence of LET-2 LLL homotrimers (Fig. 4 C).

**Figure 4.**
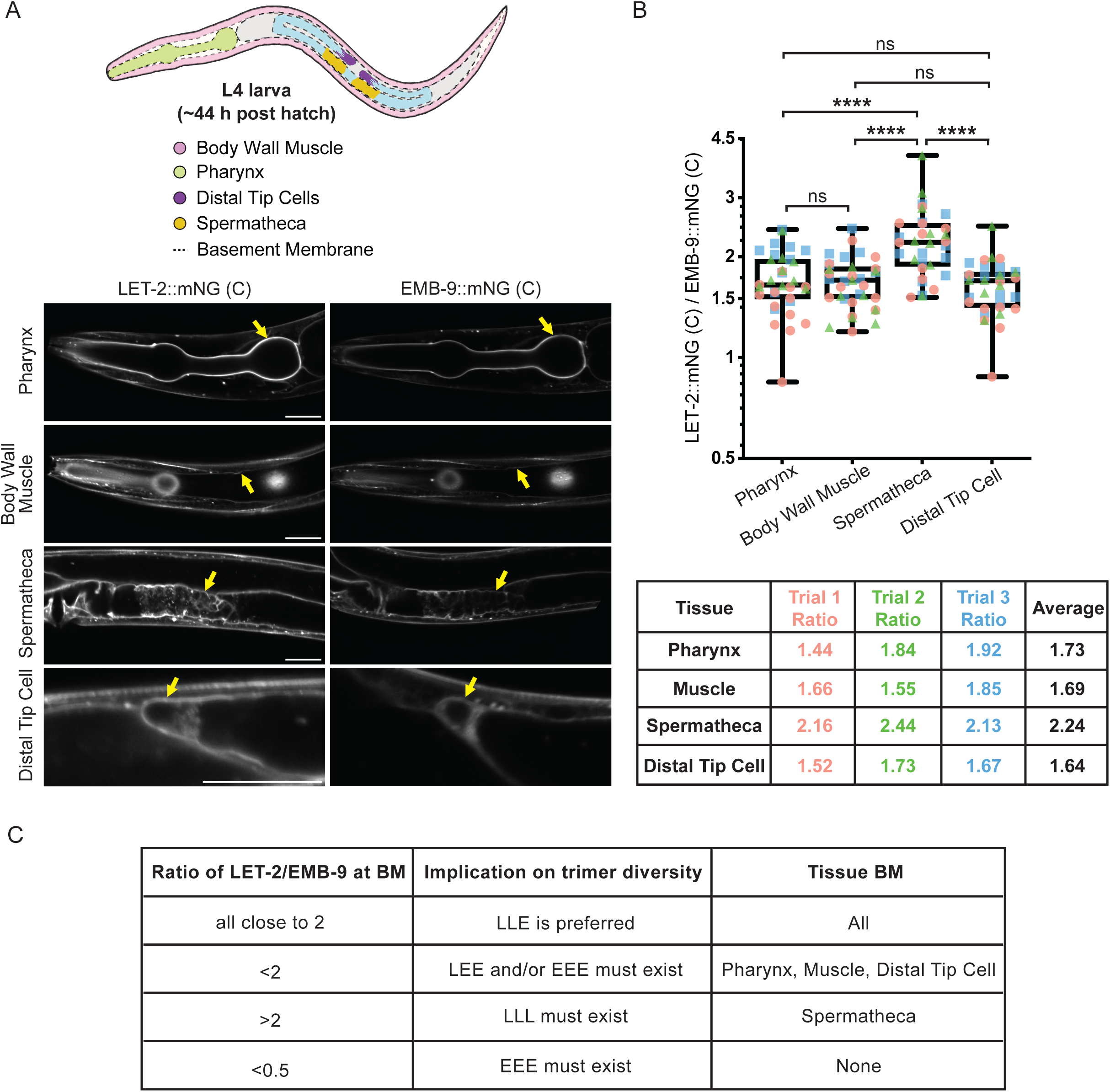
LET-2/EMB-9 BM ratios are not precisely 2:1 and differ between tissues. **(A)** Top: A schematic diagram of a late L4 larva with analyzed tissues and surrounding BMs (dashed lines) shown. Bottom: Single z-slice confocal fluorescence images showing LET-2::mNG (C) and EMB-9::mNG (C) in tissues analyzed. Yellow arrows indicate BMs measured (see Methods). The scale bar is 20 μm. **(B)** Top: Boxplot of the mean BM fluorescence intensity measurements of LET-2::mNG (C) normalized to the average of all mean BM fluorescence intensity measurements of EMB-9::mNG (C) for the same tissue and trial (see Methods, n ≥ 10 animals for each of three trials, **** P > 0.0001, ns, not significant, Brown-Forsythe ANOVA tests followed by Dunnett’s T3 multiple comparisons test). Colors of the data points indicate different trials. Bottom: Table showing the ratios of the average BM fluorescence levels of LET-2::mNG (C) / EMB-9::mNG (C) for each tissue and trial. Colors in the table correspond to data points in the Boxplot. Ratios at the spermathecal BM were consistently over 2 (LET-2::mNG (C) levels were more than double EMB-9::mNG (C) levels), while ratios at other BMs were consistently under 2. **(C)** A table showing what the composite ratio of LET-2/EMB-9 BM protein levels suggests for trimer diversity and the tissues where those ratios were found.

To confirm the validity of these measurements, we assessed both the stability of fluorophore association to EMB-9 and LET-2 C-termini and the possibility of fluorophore self-quenching, as type IV collagen networks associate end-to-end at the C-terminus (NC1 domain) possibly placing multiple fluorophores in close proximity (Bae et al., 2021). Western blot analysis using mNG antibodies confirmed that EMB-9 and LET-2 C-terminal mNG fusion proteins were present almost exclusively as full-length proteins (Fig. S4 B). To test for quenching, we reasoned that reducing the number of fluorophores by half in heterozygous animals, would reduce quenching (if present) and result in > 50% fluorescence of the homozygous strain. Fluorescence intensity measurements of EMB-9::mNG (C) and LET-2::mNG (C) L4.8 – L4.9 staged heterozygous worms, however, was not >50% of the levels of homozygous animals (Fig. S4 C), suggesting that fluorescence quenching was not occurring in homozygous animals. We conclude that mNG is stably associated with C-terminally tagged EMB-9 and LET-2 and that C-terminal inserted mNG does not show quenching in the BM. Further, the levels of LET-2/EMB-9 are close to 2:1 for all tissue BMs, consistent with a predominant LLE heterotrimer composition. However, the diversity in α-chain ratios also suggests some flexibility in the α-chain make-up of trimers.

### Tissue specific translation levels of LET-2 and EMB-9 mirror the BM α-chain ratios

We hypothesized that differences in levels of translation and thus availability of LET-2 and EMB-9 α-chains for trimerization in the ER might influence the tissue specific differences in α-chain ratios at the BM. We thus used CRISPR/Cas9 to insert a P2A peptide sequence followed by PEST::mNG immediately before the *emb-9* and *let-2* terminal codons (*emb-9::P2A::PEST::mNG* and *let-2::P2A::PEST::mNG*). The P2A peptide causes the ribosome to fail in making a peptide bond between the glycine and proline near the C-terminus of the 2A peptide during translation, resulting in the separate synthesis of EMB-9 or LET-2 and mNG::PEST from the same mRNA (Lartey et al., 2024; Sharma et al., 2012). The PEST destabilized mNG that remains within the cell cytoplasm can be used as a sensitive reporter of active EMB-9 and LET-2 translation levels (Li et al., 1998; Stevenson et al., 2023). We examined the average fluorescence intensity of PEST::mNG in the cytosol of three tissues that express and secrete type IV collagen—the body wall muscle, spermatheca myoepithelium, and the DTC (Fig. 5 A) (Graham et al., 1997). The ratio of LET-2/EMB-9 protein translation in the body wall muscle and DTC was ∼1.5 (Fig. 5 B and Fig. S4 D), similar to the ratio of the LET-2 and EMB-9 proteins in the body wall muscle and DTC BMs (Fig. 4 B and Fig. S4 A). In contrast, the ratio of LET-2/EMB-9 protein translation in the spermatheca was ∼2.5 (Fig. 5 B and Fig. S4 D), which mirrored the higher LET-2/EMB-9 protein ratio specifically found in the spermathecal BM (Fig. 4 B and Fig. S4 A). Together these findings support the idea that the levels of α-chains available for trimerization influence the α-chain composition of trimers assembled and then incorporated into BM.

**Figure 5.**
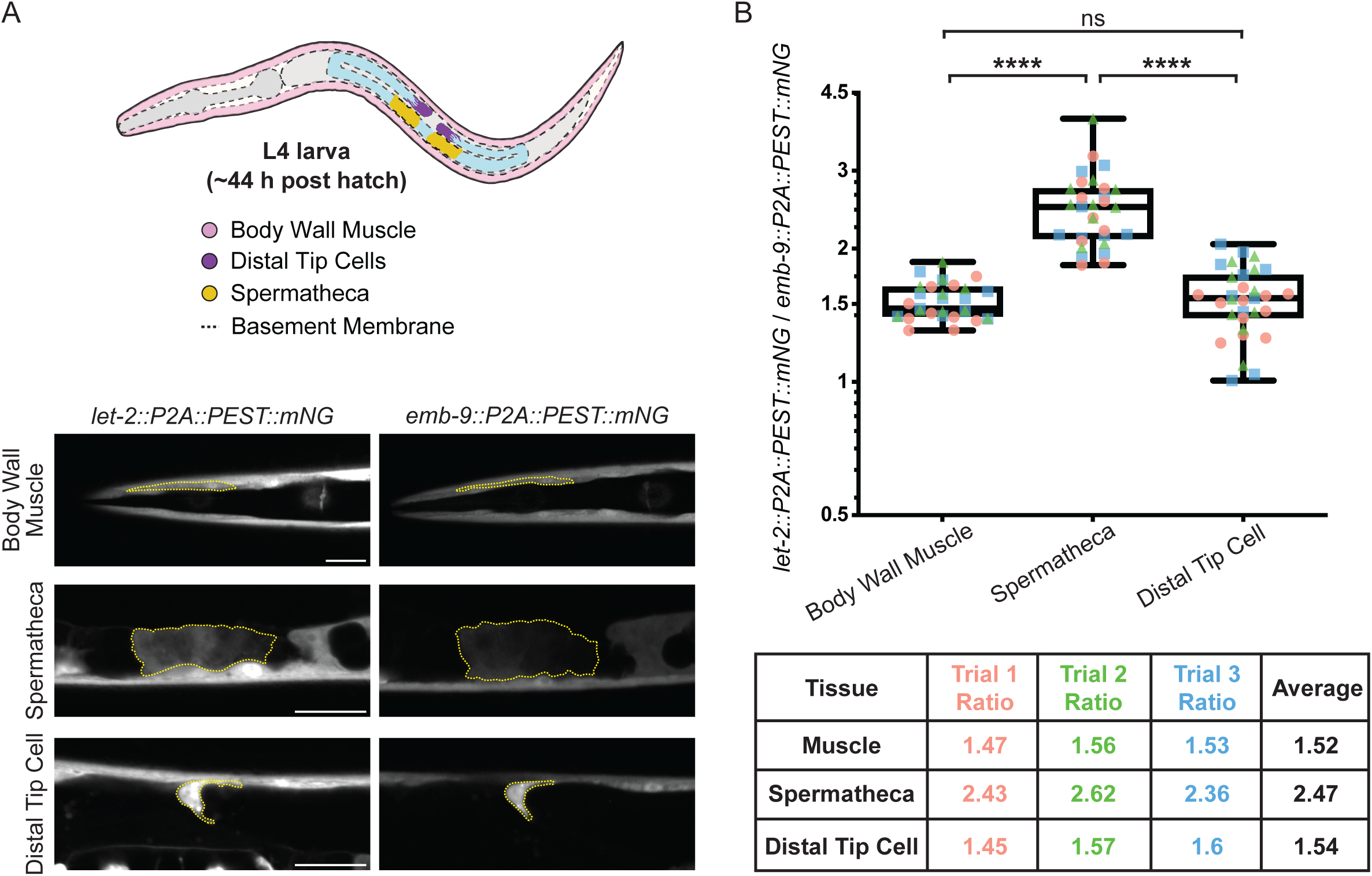
LET-2/EMB-9 translation reporter ratios in tissues mirror BM ratios. **(A)** Top: A schematic diagram of a late L4 larva with tissues analyzed for LET-2 and EMB-9 translation reporters shown. Bottom: Single z-slice confocal fluorescence images of translation reporters for LET-2 (*let-2::P2A::PEST::mNG*) and EMB-9 (*emb-9::P2A::PEST::mNG*) in the body wall muscle, spermatheca and DTC in late L4 larvae. Yellow dotted lines indicate where measurements were taken. The scale bar is 20 μm. **(B)** Top: Boxplot of the mean fluorescence intensity measurements of the LET-2 translational reporter normalized to the average of all mean fluorescence intensity measurements of the EMB-9 translational reporter in the same tissue and trial (see Methods, n ≥ 10 animals for each of three trials. **** P > 0.0001, ns, not significant, Kruskal-Wallis test followed by Dunn’s multiple comparisons test). Colors of the data points indicate different trials. Bottom: Table showing the ratios of the average fluorescence levels of LET-2 translation reporter to EMB-9 translation reporter in each trial. Colors in the table correspond to data points in the Boxplot.

### Changes in type IV collagen α-chain expression is sufficient to alter α-chain BM ratios

Our results suggest that although the ratio of LET-2/EMB-9 is close to 2:1 in tissue BMs, and thus likely predominantly contains an LLE trimer, there is also flexibility in the α-chain composition of trimers assembled and incorporated into the BM based on α-chain availability during trimer assembly. We next wanted to directly test both the preference for heterotrimer formation and if changing α-chain levels is sufficient to alter the ratios of α-chains deposited in the BM. We first reduced *emb-9* expression in EMB-9::mNG (C) and LET-2::mNG (C) knock-in worms by feeding RNAi targeting *emb-9* beginning at the L1 larval stage (Jayadev et al., 2019). We then measured the mean fluorescence intensity of EMB-9::mNG and LET-2::mNG in the body wall muscle BM of day 1 adults (72 hours post RNAi initiation). RNAi targeting of *emb-9* resulted in a 75% loss of EMB-9 protein at the body wall muscle BM (Fig. 6 A and B; Fig. S5 A). LET-2 levels in the BM were decreased by ∼50% and there was increased internal aggregation of LET-2::mNG in body wall muscle (Fig. 6 A and B; Fig. S5 A and C). The reduction in LET-2 at the BM and increase in internalization is consistent with a preference for heterotrimer assembly. Yet, a greater proportion of LET-2 than EMB-9 was incorporated into the BM—the LET-2/EMB-9 ratio went from ∼1.8 to ∼3.5 after *emb-9* knock down (Fig. 6 C and Fig. S5 B), also indicating flexibility in trimer α-chain composition based on expression levels of the α-chains.

**Figure 6.**
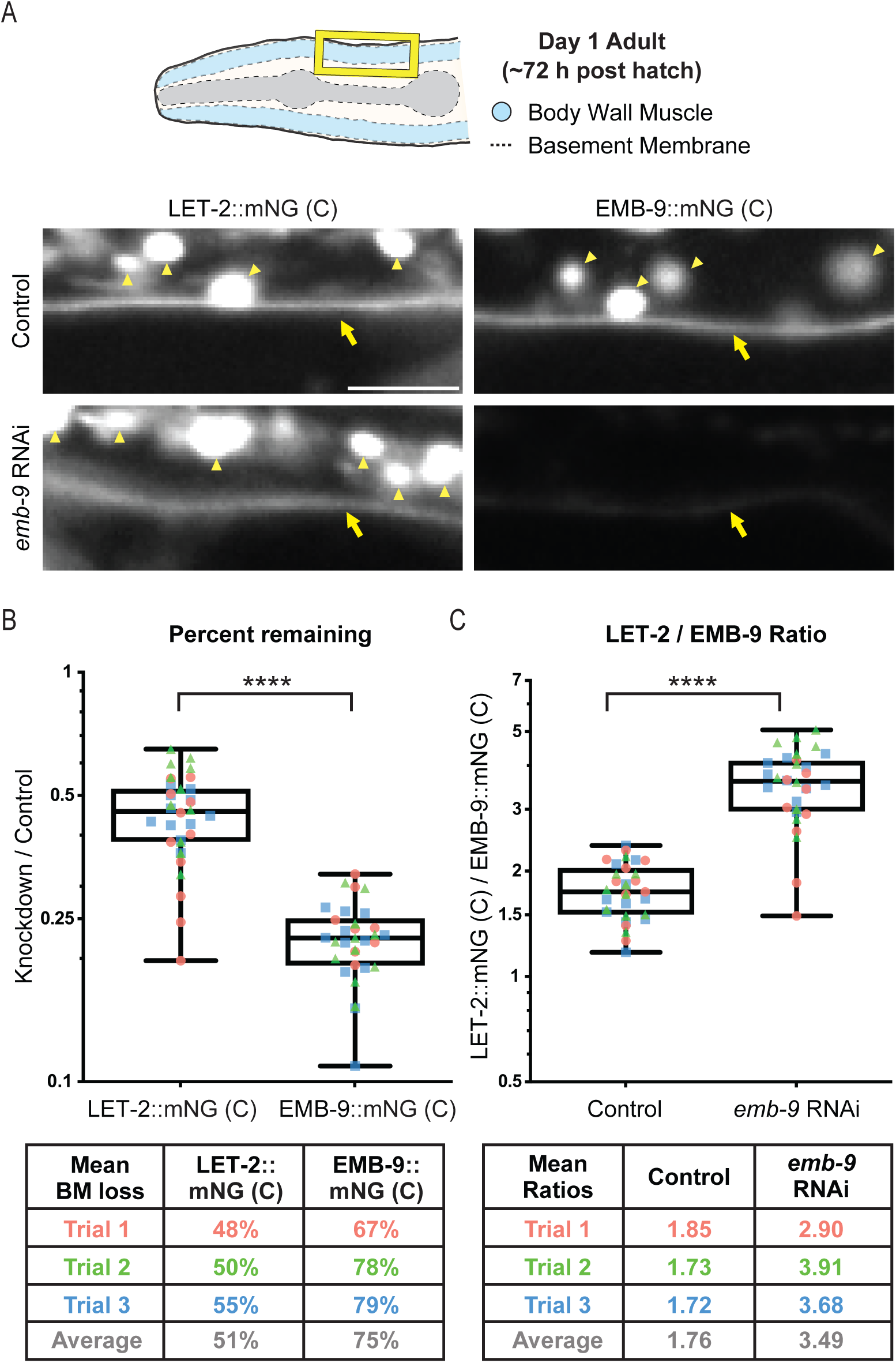
Reduction of *emb-9* expression increases the LET-2/EMB-9 BM ratio. **(A)** Top: A schematic diagram of a day 1 adult showing the head region and the location of the body wall muscle BM analyzed (yellow box). Below: Single z-slice confocal images of LET-2::mNG (C) and EMB-9::mNG (C) along the body wall muscles of day 1 adults with and without *emb-9* RNAi treatment. The yellow arrows indicate body wall muscle BM and yellow arrowheads the intracellular aggregates. Aggregates were more prevalent in LET-2::mNG (C) animals after *emb-9* RNAi treatment (see Fig. S5 C). The scale bar is 5 µm. **(B)** Top: Boxplots show percent of LET-2::mNG (C) and EMB-9::mNG (C) protein (BM mean fluorescence intensity) remaining at the BM after *emb-9* RNAi mediated knockdown. For each trial, mean fluorescence intensity measurements of LET-2::mNG (C) and EMB-9::mNG (C) with *emb-9* RNAi were normalized to the average of all mean fluorescence intensity measurements of LET-2::mNG (C) and EMB-9::mNG (C) with control RNAi (L4440 empty RNAi vector) from that trial (see Methods, n ≥ 8 animals for each of three trials. **** P > 0.0001, unpaired two-tailed Student’s t-test with Welch correction). Colors of the data points indicate different trials. Bottom: The percent loss of LET-2::mNG (C) and EMB-9::mNG (C) mean fluorescence intensity for each trial is listed in the table. Colors in the table correspond to data points in the Boxplot. **(C)** Top: Boxplots show the LET-2::mNG (C) / EMB-9::mNG (C) mean fluorescence ratios in the body wall muscle BM with *emb-9* RNAi and control RNAi (L4440 empty vector) treatment. LET-2::mNG (C) mean fluorescence intensity measurements were normalized to the average of all EMB-9::mNG (C) mean fluorescence intensity measurements within each trial (n ≥ 8 animals for each of three trials. **** P > 0.0001, unpaired two-tailed Student’s t-test with Welch correction). Bottom: Table showing the ratios of the average BM fluorescence levels of LET-2::mNG (C) / EMB-9::mNG (C) for each trial. Colors in the table correspond to data points in the Boxplot.

To complement RNAi mediated reduction of EMB-9, we also increased EMB-9 protein levels by overexpressing a tandem tagged *emb-9p*::EMB-9::mNG::mRuby2 in EMB-9::mNG (C) knock-in animals and we overexpressed *emb-9p*::EMB-9::mRuby2 in LET-2::mNG (C) knock-in animals. We used the mRuby2 tag to ensure that the overexpressed EMB-9 (marked with mRuby2) was being incorporated into the BM of these strains. Overexpression of EMB-9 increased EMB-9 levels ∼36% in the BM of day 1 adults and there was no significant change in LET-2 levels (Fig. 7 A and B; Fig. S5 D). The LET-2/EMB-9 ratio decreased from 1.7 to 1.3 (Fig. 7 C and Fig. S5 E). There was also an increase in EMB-9::mNG aggregation in the body wall muscle (but not LET-2) in animals overexpressing EMB-9 (Fig. S5 F), consistent with a preference for heterotrimer formation. Taken together, these experiments support the idea that there is a preference for LLE heterotrimer assembly, but also indicate that differences in α-chain expression can drive the flexibility in trimer α-chain make-up at the BM.

**Figure 7.**
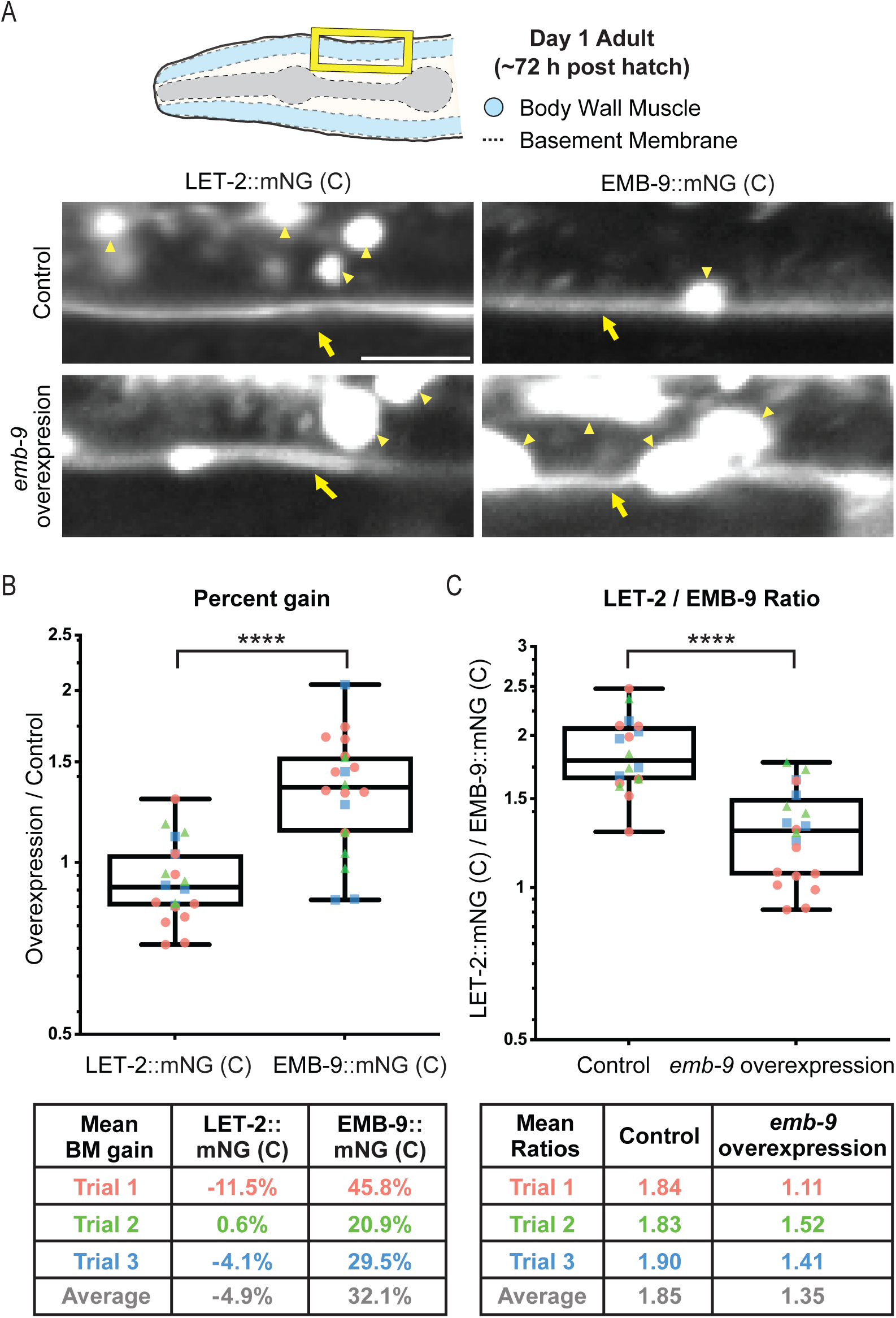
Increasing *emb-9* expression decreases the LET-2/EMB-9 BM ratio. **(A)** Top: A schematic diagram of a day 1 adult showing the head region and the location of the body wall muscle BM analyzed (yellow box). Below: LET-2::mNG (C) with and without *emb-9::mRuby2* overexpression and EMB-9::mNG (C) with and without *emb-9::mNG::mRuby2* overexpression in the body wall muscles of day 1 adults. The yellow arrows indicate body wall muscle BM and yellow arrowheads the intracellular aggregates. Aggregates were more prevalent in EMB-9::mNG (C) animals after *emb-9* overexpression (see Fig. S5 F). The scale bar is 5 µm. **(B)** Top: Boxplots show percent gain of LET-2::mNG (C) and EMB-9::mNG (C) protein (mean fluorescence intensity) in the body wall muscle BM after *emb-9* overexpression. For each trial, mean fluorescence intensity measurements of LET-2::mNG (C) and EMB-9::mNG (C) with *emb-9* overexpression were normalized to the average of all mean fluorescence intensity measurements of LET-2::mNG (C) and EMB-9::mNG (C) without overexpression (see Methods, n ≥ 5 animals for each of three trials. **** P > 0.0001, unpaired two-tailed Student’s t-test with Welch correction). Bottom: The percent gain of both proteins for each trial is listed in the table. Colors in the table correspond to data points in the Boxplot. **(C)** Top: Boxplots show the LET-2::mNG (C) / EMB-9::mNG (C) mean fluorescence ratios in the body wall muscle BM with and without *emb-9* overexpression. LET-2::mNG (C) mean fluorescence intensity measurements were normalized to the average of all EMB-9::mNG (C) mean fluorescence intensity measurements within each trial (n ≥ 5 animals for each of three trials. **** P > 0.0001, unpaired two-tailed Student’s t-test with Welch correction). Bottom: Table showing the ratios of the average BM fluorescence levels of LET-2::mNG (C) / EMB-9::mNG (C) for each trial. Colors in the table correspond to data points in the Boxplot.

### Type IV collagen glycine substitution mutant alleles cause extracellular turnover defects

Pathogenic variants in *COL4A1* and *COL4A2* lead to Gould Syndrome, a genetically dominant heterogenous multisystem disorder (Labelle-Dumais et al., 2024). The most common variants found in Gould Syndrome are substitutions in glycine amino acids in the Gly-X-Y motifs of the triple helical domain, which lead to ER retention, reduced secretion, and defective BMs in mice and humans (Gould et al., 2007; Jeanne and Gould, 2017; Kuo et al., 2014; Murray et al., 2014). Similar to humans and mice, immunofluorescence studies in *C. elegans* have revealed that *emb-9* and *let-2* glycine missense mutant alleles that mimic Gould syndrome result in intracellular accumulation and decreased BM levels (Gupta et al., 1997). Treatment of mice and human fibroblast cells with the chemical chaperone 4-phenylbutyrate (4PBA) increases secretion of mutant type IV collagen and increases incorporation into mice BMs, but does not alleviate disease severity for all tissues, and can even increase skeletal myopathy for some *COL4A1* mutant alleles (Gould et al., 2007; Jones et al., 2019; Kuo et al., 2014; Labelle-Dumais et al., 2019). This suggests that type IV collagen harboring glycine substitutions might not function normally extracellularly. How these mutant forms may alter type IV collagen extracellular regulation and function, however, is unknown.

We next wanted to assess whether the ability to endogenously visualize fluorophore tagged type IV collagen with quantitative live imaging could reveal new insights into how glycine substitutions affect type IV collagen extracellular regulation. Previous immunolocalization studies in C. *elegans* glycine substitution mutants in *emb-9* and *let-2* have reported apparent decreased type IV collagen α-chain BM levels, but these levels were not quantified (Gupta et al., 1997). We thus used genome editing to tag the temperature sensitive glycine substitution mutants *emb-9* (*b117)* (EMB-9 G1173D) and *let-2* (*b246)* (LET-2 G1287D) with mNG at the C-terminus (Fig. 8 A) (Gupta et al., 1997; Sibley et al., 1994). Animals harboring *emb-9* (*b117)* or *let-2* (*b246)* are homozygous viable at the permissive temperature of 15°C but mothers shifted to 25°C have progeny that die as embryos or arrest as larvae (Gupta et al., 1997; Sibley et al., 1994).

**Figure 8.**
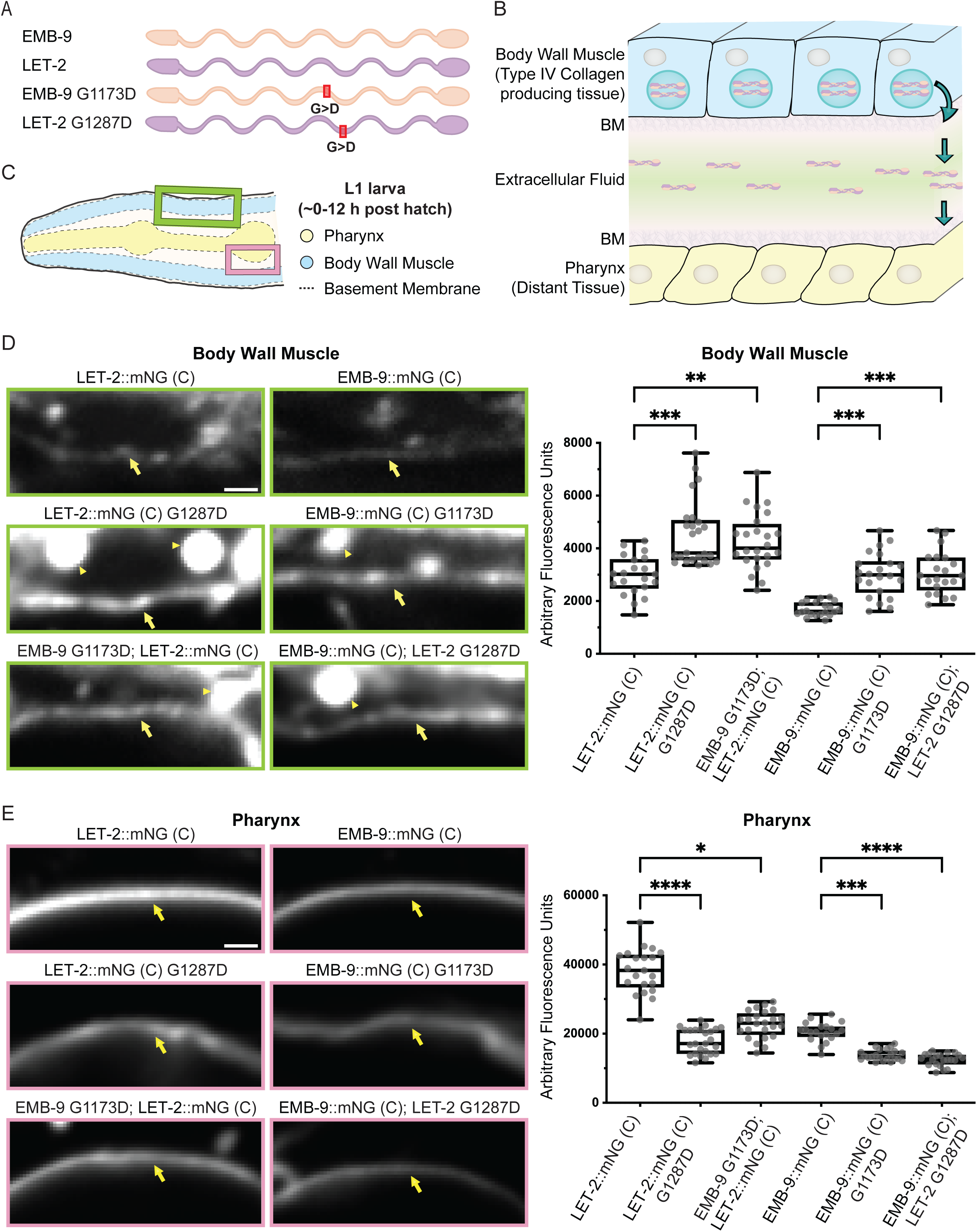
EMB-9 and LET-2 glycine substitution mutants mimicking Gould syndrome accumulate at higher levels in body wall muscle BM. **(A)** A schematic indicating the location of glycine to aspartic acid mutations in EMB-9 and LET-2 proteins. **(B)** A schematic of the *C. elegans* body wall muscles, which are the primary site of type IV collagen production. Secreted type IV collagen is incorporated into body wall muscle BM or trafficked in extracellular fluid and incorporated into the BM of distant tissues (such as the pharyngeal BM). **(C)** A schematic showing BM locations analyzed in L1 larvae. The green box corresponds to images in Fig. 8 D and the pink box corresponds to images in Fig. 8 E. **(D)** Left: Single z-slice confocal fluorescence images in the body wall muscles of LET-2::mNG (C) and EMB-9::mNG wild-type animals, LET-2::mNG G1287D and EMB-9::mNG (C) G1173D mutant animals, and animals with a wild-type fluorescently tagged protein paired with an untagged mutant protein: EMB-9 G1173D; LET-2::mNG (C) and EMB-9::mNG (C); LET-2 G1173D in the body wall muscles. Yellow arrows indicate body wall muscle BM and yellow arrowheads indicate aggregates in body wall muscle cells (see Fig. S5 G). Right: Boxplots showing mean fluorescence intensity measurements of body wall muscle BM (n ≥ 10 animals per condition**, \*\***P < 0.01, ***P < 0.001, Kruskal Wallis followed by Dunn’s multiple comparison test) **(E)** Left: Single z-slice confocal fluorescence images in pharynx of LET-2::mNG (C) and EMB-9::mNG (C) wild-type animals, LET-2::mNG (C) G1287D and EMB-9::mNG (C) G1173D mutant animals, and animals with a wild-type fluorescently tagged protein paired with an untagged mutant protein: EMB-9 G1173D; LET-2::mNG (C) and EMB-9::mNG (C); LET-2 G1173D. Yellow arrows indicate body wall muscle BM (note lack of intracellular aggregates in the pharynx, which does not express type IV collagen). Right: Boxplots showing mean fluorescence intensity measurements of pharyngeal BM (n ≥ 10 animals per condition**, \***P < 0.1, ***P < 0.001, ****P < 0.0001, Kruskal Wallis followed by Dunn’s multiple comparison test). The scale bars are 1 μm and all animals were grown at 25°C (restrictive temperature for *emb-9* and *let-2* mutants).

We examined type IV collagen localization at the body wall muscle BM, where most type IV collagen is generated and secreted, and at the pharynx—a tissue that does not produce type IV collagen, but instead recruits it from extracellular fluid (Fig. 8 B and C) (Graham et al., 1997; Morrissey et al., 2016). We examined mNG (C) tagged EMB-9 G1173D and LET-2 G1287D mutants arrested at the L1 larval stage and compared localization to mNG (C) tagged EMB-9 and LET-2 wild-type L1 larvae. In line with previous studies, we saw extensive intracellular accumulation in mutant strains at non-permissive temperature (Fig. S5 G). Interestingly, we found that the levels of mutant EMB-9 (G1173D) and LET-2 (G1287D) protein were higher in the body wall muscle BM compared to non-mutant control proteins, but the mutant protein levels were lower in the pharyngeal BM (Fig. 8 D and E). Consistent with a preference for heterotrimers, we found that when wild-type mNG tagged EMB-9 and LET-2 α-chains were crossed into the respective untagged *let-2* (*b246)* and *emb-9* (*b117)* mutant backgrounds, we observed similar intracellular accumulation, but also increased body wall muscle BM levels and decreased pharyngeal BM levels (Fig. 8 D and E; Fig. S5 G). As type IV collagen is trafficked from the body wall muscles to other tissues such as the pharynx, these results suggest that mutant type IV collagen might not be properly removed from the body wall muscle BM to allow for its distribution to other tissues.

To examine removal dynamics of mutant type IV collagen trimers from the body wall muscle BM, we crossed animals with endogenously tagged photoconvertible EMB-9::mEos2 and LET-2::mMaple into untagged *let-2*(*b246)* and *emb-9*(*b117)* mutants, respectively. We examined the removal of type IV collagen from the body wall muscle BM in wild type and mutant animals by photoconverting the body wall muscle BM and assessing reduction in photoconverted signal after five hours. Approximately 50% of photoconverted signal was removed in both EMB-9 (G1173D) and LET-2 (G1287D) mutants after five hours, while 70% was removed in wild-type EMB-9 and LET-2 controls (Fig. 9 A-C). Together these results indicate that glycine missense mutant α-chains cause BM accumulation defects that might result from decreased removal from BMs surrounding tissues that produce type IV collagen.

**Figure 9.**
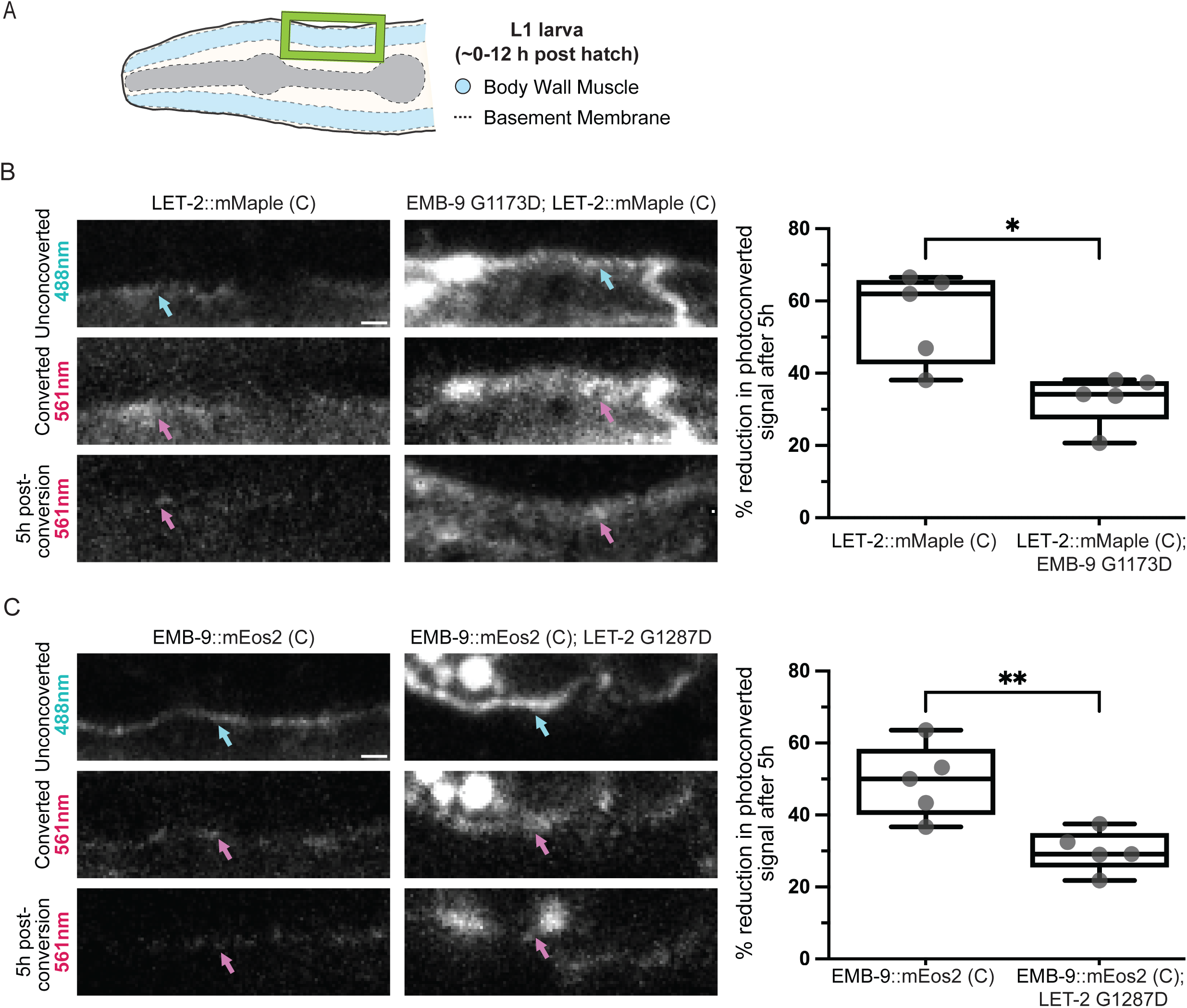
EMB-9 and LET-2 glycine substitution mutants have slower removal rates from body wall muscle BM. **(A)** The schematic indicates body wall muscle BM region analyzed (yellow box) in L1 larvae. **(B)** Left: Single z-slice representative confocal fluorescence images of the LET-2::mMaple (C) signal at the body wall muscle BM in wild-type animals (LET-2::mMaple (C)) or *emb-9* glycine mutant animals (EMB-9 G1173D; LET-2::mMaple (C)). Unconverted fluorophore signal (488 nm excitation), converted fluorophore signal (561 excitation) immediately after photoconversion, and 5 h post-photoconversion (561 excitation) are shown. Cyan and magenta arrows point to unconverted and converted signal at the body wall muscle BM, respectively. Right: Boxplots show the percent of photoconverted signal reduced after 5 h (n ≥ 5 animals, P > 0.05, unpaired two-tailed Student’s t-test). **(C)** Left: Single z-slice representative confocal fluorescence images of the EMB-9::mEos2 (C) signal at the body wall muscle BM in wild-type animals (EMB-9::mEos2 (C)) or *let-2* glycine mutant animals (EMB-9::mEos2; LET-2 G1287D). Unconverted fluorophore signal (488 nm excitation), converted (561 excitation) immediately after photoconversion, and 5 h post-photoconversion (561 excitation) are shown. Cyan and magenta arrows point to unconverted and converted signal at the body wall muscle BM, respectively. Right: Boxplots show the percent of photoconverted signal reduced after 5 h (n ≥ 5 animals, P > 0.05, unpaired two-tailed Student’s t-test followed by Mann-Whitney test). All animals were grown and recovered at 25°C (restrictive temperature for *emb-9* and *let-2* mutants). The scale bars are 1 μm.

## Discussion

Type IV collagen is built as a helical trimer, composed of three interweaved α-chains. Comparative studies in animals suggest that type IV collagen is a founding and core BM component with crucial structural and signaling roles throughout animal life (Fidler et al., 2018; Poschl et al., 2004). Most vertebrates encode six α-chains, while *C. elegans* and *Drosophila* each have two (Fidler et al., 2017). Genetic and regulatory defects in type IV collagen are implicated in numerous human diseases and aging (Gatseva et al., 2019; Mao et al., 2015; Uspenskaia et al., 2004; Zhang et al., 2022). Thus, there is broad interest in understanding type IV collagen regulation and function. Endogenous fluorophore tagging has powerfully advanced the understanding of BM protein regulation and function in *Drosophila* and *C. elegans*, and more recently in zebrafish and mice (Stramer and Sherwood, 2024). Endogenous tagging of type IV collagen α-chains in *Drosophila*, *C. elegans*, and mice has almost exclusively used the *Drosophila* Vkg-GFP protein trap insertion site, where GFP is inserted at the start of the 7S domain near the N-terminus of the protein (Jones et al., 2024; Keeley et al., 2020; Morin et al., 2001; Stramer and Sherwood, 2024). Insertion of fluorophores at this site is viable for *Drosophila vkg*, *C. elegans emb-9*, and mouse *COL4A2* α-chains (Keeley et al., 2020; Morin et al., 2001; Wuergezhen et al., 2025), but has not yet been attempted for *Drosophila Cg25c*, and is not viable for *C. elegans let-2* or mouse *COL4A1* α-chains (Jones et al., 2024; Keeley et al., 2020). As a result, there has been an inability to comprehensively examine type IV collagen regulation and composition using live imaging in any animal to date.

Using a health assay and localization analysis, we found that *C. elegans emb-9* tagged at the Vkg-GFP site, which we have named internal site 1 (IS1), causes reduced health and extensive aggregation of fluorophore tagged protein in the ER of body wall muscle cells, where type IV collagen is predominantly expressed and secreted (Graham et al., 1997). Tagging *emb-9* at the IS1 site also reduced the amount of EMB-9 protein localized to BM. We also found that *emb-9* tagged at an internal site within an interruption of the Gly X–Y repeats (IS2) (Matsuo et al., 2019), had reduced health and had ER localized aggregation of tagged EMB-9 protein. In contrast, we discovered that tagging the *C. elegans* type IV collagen α-chains EMB-9 and LET-2 at the C-terminus with a linker and many different fluorophores resulted in animals that were indistinguishable from wild-type animals in our health assay. Even animals that were homozygous for both labelled α-chains (*e.g.,* EMB-9::mRuby2 (C); LET-2::mNG (C)) were similar to wild-type animals. Further, these type IV collagen strains showed little to no internal protein aggregation in early larval staged worms. Western blot analysis also confirmed that the fluorophore mNG was stably associated with both EMB-9 and LET-2. One reason that the C-terminus of type IV collagen α-chains might not have been selected for endogenous tagging previously, is that the NC1 domain, which both selects for heterotrimer assembly and mediates dimerization of type IV collagen molecules extracellularly through sulfilimine bonds, is located at the C-terminus (Fidler et al., 2018; Khoshnoodi et al., 2008). C-terminally tagged type IV collagen trimers could also cluster multiple fluorophores-up to six when both α-chains are tagged. Yet, we did not detect fluorophore quenching (Bae et al., 2021), suggesting that C-terminal site tagging, along with the 18 amino acid linker we used, does not cause quenching in native BM.

One unexpected finding from our studies was the dramatic influence of different fluorophores inserted at the IS1 (Vkg) site on strain health. Insertion of the red fluorophores mRuby2 or mScarlet-I resulted in only mild delays in the time to starvation health assay for worm populations (1 day delay). In contrast, insertion of mEos2 or mNG, resulted in dramatically slower growth and long delays to starvation (5-8 days). All these genetically encoded fluorescent proteins are similar in size, share the characteristic β-barrel structure of the Aequorea GFP, and have similar mechanisms of fluorophore formation (Rodriguez et al., 2017). However, red fluorescent proteins and photoconvertible proteins have been constructed through extensive engineering of genes from a variety of anthozoan species and are thus encoded by significantly different amino acids (Bindels et al., 2017; Kredel et al., 2009; Thorn, 2017; Zhang et al., 2012). As a result, the folding rates and fluorophore formation rates vary significantly between fluorescent proteins (Balleza et al., 2018; Thorn, 2017). Our results suggest that these differences might impact the ability of type IV collagen α-chains to fold and trimerize in the ER, particularly when tagged at the IS1 (Vkg) site, and negatively affect animal health.

Using our endogenous C-terminal fluorophore tagged LET-2 and EMB-9 α-chains, our proteomics findings indicated that LET-2 is approximately two-fold more abundant than EMB-9 in whole animals. This observation suggests that the *C. elegans* type IV collagen trimer is composed predominantly of two LET-2 (α2-like chain) and one EMB-9 (α1-like chain)—an LLE heterotrimer composition (Guo and Kramer, 1989). Notably, this composition does not follow the proposed mammalian rule of combining two odd-numbered (α1-like) and one (α2-like) even α-chain (Khoshnoodi et al., 2008). *Drosophila* also appears to not follow the mammalian rule. Hexamer isolation, NC1 recombination experiments, and transgenic NC1 domain expression suggests that the *Drosophila* type IV collagen trimer is predominantly Cg25c-Cg25c-Vkg (*i.e.,* two α2 (Cg25c) and one α1 (Vkg) chain). This discrepancy, however, might simply be due to the difficulty of assigning clear homology of *C. elegans* and *Drosophila* NC1 sequences into vertebrate α1- and α2-like groups (Guo and Kramer, 1989; Summers et al., 2023).

A critical step in type IV collagen trimer maturation is catalyzed by prolyl 4-hydroxylase. In the absence of prolyl 4-hydroxylase or its cofactor ascorbate/vitamin C, type IV collagen α-chains cannot form trimers (Lamande and Bateman, 1999). Work in *Drosophila* using GFP tagged Vkg protein showed that in prolyl 4-hydroxylase deficient larvae, type IV collagen is not incorporated into BMs (Pastor-Pareja and Xu, 2011). These results strongly suggest that all type IV collagen within BMs are trimers, and that fluorescence intensity of individual α-chains tagged with the same fluorophore at the same site can be used to assess the composite picture of BM trimers within the BM. Importantly, our quantitative fluorescence analysis of tissue specific BMs revealed BMs did not have a strict LET-2/EMB-9 ratio of 2.0 (*i.e.,* are not stringently LLE), instead most were ∼1.7, whereas the spermathecal BM had a ratio of ∼2.2. These observations suggest a likely predominance of the LLE trimer, but also some flexibility in trimer composition. LET-2/EMB-9 ratios greater than 2 suggest the existence of LLL homotrimers, while ratios less than 2 indicate LEE and possibly EEE trimers. Analysis of translational reporters within specific tissues and experimentally increasing and decreasing α-chain expression, showed that trimer compositional variability within BMs can be driven by differences in α-chain expression levels. This implies that the relative availability of α-chains for trimerization in the ER influences trimer make-up. Notably, our studies also revealed a dependence between α-chains for secretion— decreasing the levels of EMB-9 led to retention of LET-2 in cells, while increasing EMB-9 expression led to EMB-9 intracellular accumulation. This observation is consistent with studies showing that mutations in *emb-9* and *let-2* result in intracellular retention of the other (non-mutant) α-chain (Gupta et al., 1997). Together, these results suggest a preference for the in vivo assembly of the LLE heterotrimer, but also some flexibility in building other trimers in *C. elegans*.

Based on α-chain abundance and hexamer associations of the six α-chains in vertebrates, it’s been proposed that only three (out of a possible 56) type IV collagens trimers are built--α1α1α2, α3α4α5, and α5α5α6 (Khoshnoodi et al., 2008). Yet, studies on the α1α1α2 trimer, the most prevalent and most investigated type IV collagen in vertebrates, suggest that other trimer combinations could exist--carcinoma cells secrete α1 homotrimers (Haralson et al., 1985), reconstitution of NC1 domains have revealed many possible trimer combinations (Boutaud et al., 2000), and in vivo hexamer analysis has not excluded the existence of α1 homotrimers or α1α2α2 heterotrimers (Boutaud et al., 2000; Johansson et al., 1992; Siebold et al., 1988). Studies in *Drosophila* have also not ruled variable trimer formation (Summers et al., 2023). Thus, flexible type IV collagen trimer combinations might occur in other organisms.

Trimer compositional variability likely expands the possible functions of type IV collagen and could allow for tissue-specific tailoring of type IV collagen trimer construction. For example, it’s been hypothesized that the more restricted and tissue specific vertebrate α3α4α5 heterotrimer might have increased mechanical strength relative to that of the α1α1α2 heterotrimer because of additional internal cross-linking (Gunwar et al., 1998; Naylor et al., 2021). Interestingly, we found that the spermathecal BM had a LET-2/EMB-9 ratio of ∼2.2, whereas other tissue BMs had a significantly lower ratio of ∼1.7. The spermatheca tissue is unique in that it undergoes repeated stretching every ∼20 minutes during peak egg laying in *C. elegans* (Kelley and Cram, 2019). The distinct type IV collagen composition found in the spermatheca BM might mechanically support the ability of the spermatheca to repetitively stretch. Trimer α-chain compositional diversity could also increase the variety and specificity of interactions with other BM components, signaling molecules, and cell surface receptors (Khoshnoodi et al., 2008; Wang et al., 2008) to support many tissue-specific functions.

The most common human pathogenetic variants in the *COL4A1* and *COL4A2* genes are those that result in substitutions of glycine residues with other amino acids within the triple helical domain (Jeanne and Gould, 2017). These variants often impair the formation and secretion of COL4A1 and COL4A2 heterotrimers (α1α1α2) and lead to Gould syndrome, which is a multisystem disorder involving variable defects in many tissues, including cerebrovascular, eye, kidney, brain, skeletal, and neuromuscular (Labelle-Dumais et al., 2024; Mao et al., 2015). Notably, not all glycine mutations affect type IV collagen secretion (Kuo et al., 2014). Further, pharmacological treatment of mice harboring *COL4A1* mutations with the chemical chaperone 4-phenylbutyrate (4PBA), which enhances secretion and BM incorporation of mutant forms of type IV collagen, can decrease, increase, or have no effect on tissue pathology depending upon the mutation and tissue (Hayashi et al., 2018; Jones et al., 2019; Labelle-Dumais et al., 2019). These observations highlight the complex roles of type IV collagen in tissue function and suggest that mutations in type IV collagen can affect its extracellular functions. Through C-terminal tagging with mNG, we show that temperature sensitive mutants harboring glycine substitutions in *emb-9* and *let-2* can be successfully labelled with fluorophores. We found that mutant forms of both α-chains led to type IV collagen retention in cells at the non-permissive temperature, as previously reported using immunofluorescence (Gupta et al., 1997). However, we also discovered increased accumulation of mutant type IV collagen in the BM on the body wall muscle, which is a site of type IV collagen production and secretion, and decreased accumulation of type IV collagen on the pharynx—a tissue that does not produce type IV collagen (Graham et al., 1997). Through optical highlighting of photoconvertible mEos2 and mMaple tagged type IV collagen trimers, we found that type IV collagen harboring glycine substitution mutations were more stably associated with the body wall muscle BM. This decrease in type IV collagen BM on-off rate, might not only increase accumulation in BMs surrounding cells and tissues that secrete type IV collagen, but also decrease levels in BMs surrounding tissues such as the pharynx, that depend upon type IV collagen trafficking away from secreting tissues (Duncan et al., 2020; Pastor-Pareja and Xu, 2011; Stramer and Sherwood, 2024). Consistent with this notion, a similar increase of type IV collagen in body wall muscle BM and decrease in pharyngeal BM occurs after loss of the extracellular collagen chaperone SPARC, which promotes type IV collagen trafficking from sites of synthesis (Morrissey et al., 2016; Pastor-Pareja and Xu, 2011). As type IV collagen levels can affect BM biomechanical and signaling properties (Crest et al., 2017; Jayadev and Sherwood, 2017b; Tian and Jiang, 2014), these alterations in type IV collagen BM accumulation could in part explain extracellular dysfunction of *COL4A1* and *COL4A2* variants.

Collagens are a major ECM superfamily composed of 46 α-chains that form 28 different collagens in mammals (I–XXVIII). All collagens share a triple helical domain, but they take on many forms, ranging from fibrils, to networks, and even membrane spanning proteins, which together play essential roles in cell adhesion, mechanical support, and signaling (Ricard-Blum, 2011; Wakabayashi, 2020). Variants in collagens are implicated in nearly 40 human genetic diseases, most of which are glycine substitutions in the triple helical domain (Fidler et al., 2018). Although generally viewed as stable, static scaffoldings, work in *Drosophila* and *C. elegans* have revealed that type IV collagen has dynamic properties, including rapid polymerization, remodeling, and association and disassociation from BM (Serna-Morales et al., 2023; Stramer and Sherwood, 2024). We expect that the tagging at the C-terminus of mature α-chains (either extreme C-terminus or prior to cleaved pro-peptide), will be broadly applicable to many collagens. Supporting this notion, type XVIII collagen, the membrane spanning COL-99, and the BLI-2 and BLI-6 cuticular collagens in *C. elegans* have been endogenously tagged with mNG at the C-terminus and are homozygous viable (Adams et al., 2023; Jayadev et al., 2022; Keeley et al., 2020). Endogenously tagging diverse collagen molecules with genetically encoded fluorophores offers to not only provide important insight into human disease, but also into the fascinating functions and likely underappreciated dynamics of this crucial ECM superfamily.

## Materials and Methods

### *C. elegans* strains and growth conditions

*C. elegans* were maintained according to standard procedures on nematode growth medium (NGM) agar plates seeded with OP50 *E. coli* at 16°C or 20°C (Stiernagle, 2006). Strains used in this study are listed in Table S1.

### DNA Cloning

#### Endogenous knock-in constructs

All genome edited strains were generated using CRISPR/Cas-9 mediated homologous recombination with a self-excising cassette (SEC) for drug selection (hygromycin) as previously described (Dickinson and Goldstein, 2016; Keeley et al., 2020). For all C-terminally tagged strains, we placed an 18 amino acid flexible linker in frame and directly upstream of the fluorophore. The flexible linker and fluorophore were inserted at the endogenous locus, just upstream of the stop codon. For *emb-9::mScarlet-I*, flexible linkers were placed on both sides of the fluorophore and the construct was inserted in frame at the same location as previously published IS1 constructs (Jayadev et al., 2022; Keeley et al., 2020). Primers used in CRISPR repair plasmid construction are listed in Table S2. We generated guide RNA (sgRNA) plasmids by inserting the sequences into the pDD122 plasmid (Table S3) (Dickinson et al., 2013). Nucleic acid sequences of sgRNAs and amino acid sequences of flexible linkers are listed in Supplementary Figure 1.

#### Translational Reporter Constructs

A *C. elegans* codon optimized P2A sequence was built using primers listed in Table S2. The PEST sequence was amplified from plasmid pCFJ150 (Table S3) (Kaymak et al., 2016) using primers listed Table S2. *let-2*::mNG (C) and *emb-9*::mNG (C) CRISPR repair plasmids were linearized by ClaI digestion. The P2A and PEST sequences were assembled with both linearized CRISPR plasmids by Gibson Assembly. C-site sgRNAs of *emb-9* or *let-2* were used to direct Cas9 cleavage for homologous insertion.

#### Transcriptional Reporter Construct

The ∼2kb *let-2* upstream regulatory sequence was amplified from wild type genomic DNA using the primers listed in Table S2. The mNG sequence was amplified from the *emb-9*::mNG (C) CRISPR repair plasmid. Both fragments were ligated into the pAP088 starter repair plasmid (Pani and Goldstein, 2018) containing homology arms for the ttTi4348 site using Gibson assembly. This single-copy transgene was inserted into the ttTi4348 transposon insertion site on chromosome I (Frokjaer-Jensen et al., 2012) using the SEC CRISPR repair plasmid as described above. The pCFJ352 plasmid (Table S3) containing the sgRNA was used to direct Cas9 cleavage near this region for insertion.

#### Overexpression Constructs

The *emb-9* overexpression construct *emb-9p::emb-9::mRuby2::emb-9 3’utr* (pQD03) was built by Gibson assembling by ligating the following fragments into pBluescript SK(-): the ∼3.8kb *emb-9* upstream regulatory sequence, the full length *emb-9* gene, an ∼500bp *emb-9* 3’utr (all amplified from N2 genomic DNA sequence) and mRuby2 amplified from the *emb-9*::mRuby2(C) CRISPR repair plasmid (using primers listed in Table S2). To create the *emb-9p::emb-9::mNG::mRuby2::emb-9 3’utr* (pQD04) construct, the mNG sequence amplified from the *emb-9*::mNG (C) CRISPR repair plasmid was inserted into pQD03 using Gibson Assembly.

### Creation of transgenic strains

Mixtures of sgRNA plasmid (5-10 ng/μL), SEC repair template plasmid (100 ng/μL), and co-injection marker plasmids (pCFJ90 [*myo-2p::mCherry*] and pCFJ104 [*myo-3p::mCherry*], 5 ng/μL each) (Table S3) were injected into the gonads of young adult N2 hermaphrodites as previously described (Dickinson and Goldstein, 2016). mNG knock-in strains harboring glycine mutation alleles for *emb-9* and *let-2* were generated using the respective C-site mNG CRISPR repair plasmid and C-site sgRNA sequence (described in *<u>Endogenous knock-in constructs</u>*). Translational reporters for *emb-9* and *let-2* were generated using the respective C-site sgRNA sequence (described in *<u>Endogenous knock-in constructs</u>*). The *let-2* transcriptional reporter was generated using the sgRNA sequence in pCFJ352 plasmid (Table S3) that targets the ttTi4348 transposon insertion site on chromosome I. Following injection, animals were singled onto OP50 plates and allowed to lay eggs at 20°C for 2-3 days before treating with 500 μL of hygromycin (2 mg/mL). After 3-4 days at 20°C post hygromycin treatment, surviving candidate knock-in animals were identified by the dominant roller phenotype (*sqt-1(e1350))* and an absence of co-injection marker fluorescence. To excise the SEC from homozygous viable lines, 4-8 L3/L4 animals were heat shocked in a 34°C water bath for 4 hrs, and then left to grow at 25°C for 3-4 days. Progeny lacking the roller phenotype were singled and edited lines were identified by visualizing the fluorophore at the BM and via PCR genotyping DNA using the primers listed in Table S2.

#### emb-9 overexpression strains

Transgenic worms expressing *emb-9p::emb-9::mRuby2::emb-9 3′utr* or *emb-9p::emb-9::mNG::mRuby2::emb-9 3′utr* were created by injecting pQD03 or pQD04 (200 ng/µl) with the plasmid pRF4 containing *rol-6(su1006)* that causes a dominant “roller” phenotype (50 ng/µl), EcoRI-digested salmon sperm DNA (30 ng/µl), and the empty pBluescript II vector (40 ng/µl) into the gonads of young adult LET-2::mNG or EMB-9::mNG hermaphrodites, respectively (Yochem and Herman, 2003). The pQD03 and pQD04 were expressed as extrachromosomal arrays, which often have mosaic somatic expression (Yochem and Herman, 2003). mRuby2 fluorescence was used to confirm transgene expression and BM localization of the overexpressed protein. The dominant roller phenotype was used to track transgene propagation. Young adult rollers (48 h post hatch) were imaged for the experiments.

### RNAi

RNAi experiments were performed using the feeding method (Timmons and Fire, 1998) and the *emb-9* RNAi construct was obtained from the Ahringer library (Fraser et al., 2000; Kamath et al., 2003). RNAi bacterial cultures were grown in selective media at 37°C for 18 hrs and then for one additional hour after adding IPTG (1 mM final concentration) (Table S3) to induce double-stranded RNA expression. Cultures were then used to seed NGM plates that had been treated with a 1:1 mixture of 1M IPTG and 100 mg/mL ampicillin (18 μL per plate) (Table S3). Plates were allowed to dry overnight after seeding. Synchronized L1 larvae were placed on the seeded plates and allowed to grow for 72 hrs at 20°C. The L4440 empty vector was used as a control (Timmons and Fire, 1998).

### Image acquisition

#### Confocal fluorescence imaging

Confocal images (excluding photoconversion experiments) were acquired at 20°C on Zeiss Axio Imager A1 microscopes, fitted with either a Yokogawa CSU-10 or Yokogawa CSU-W1 spinning disc, and either an ORCA-Fusion CMOS or ORCA-Quest qCMOS camera using 488 nm, 505 nm, and 561 nm excitation lasers. All microscopes were controlled with μManager software v1.4.23 or v2.0.1 (Edelstein et al., 2010). Zeiss 40x, 63x and 100x Plan Apochromat (1.4 numerical aperture) oil immersion objectives were used. Animals were mounted on 5% noble agar pads with 0.002M sodium azide (Table S3) for L1s and 0.01 M sodium azide for all other stages (Kelley et al., 2017).

#### Fluorescence recovery after photobleaching (FRAP) experiments

For FRAP experiments, photobleaching was performed using an iLas^2^ targeted laser system from BioVision equipped with an Omicron Lux 60mW 405nm continuous wave laser and controlled with MetaMorph version 7.10.4.407 software (Keeley et al., 2020). Laser power and duration of photobleach were consistent for all animals within an experiment. Imaging intervals and duration allowed for fluorescence recovery were determined empirically to capture quantifiable recovery. For all FRAP experiments, worms were immobilized on 5% agar pads with 0.002M sodium azide. The half of the pharyngeal terminal bulb opposite to the L1 gonad was photobleached. Animals were then rescued from the slide and washed with M9 to dilute the sodium azide treatment. Recovery was assessed using the same imaging settings after allowing the worms to develop normally for 5 hrs. Animals rescued after posterior pharynx photobleaching all developed normally and had no pharyngeal morphological defects (n = 30/30 animals).

#### Optical highlighting/photoconversion experiments

Photoconversion experiments were performed at 20°C on an inverted Zeiss 880 LSM confocal mounted to a Zeiss Axio Observer Z1 microscope using a Zeiss 63x Plan Apochromat (1.4 numerical aperture) oil immersion objective and a pinhole size of 57 μm. The photoconvertible mMaple and mEos2 fluorophores were photoconverted with a 405 nm laser and photoconversion settings were the same between samples. Unconverted mMaple and mEos2 were imaged using a 488 nm laser and converted mMaple and mEos2 with a 561 nm laser. For all optical highlighting experiments, worms were immobilized on 5% agar pads with 0.002 M sodium azide. An approximately 350 μm^2^ section of the body wall muscle BM EMB-9::mEOS2 or LET-2::mMaple between the pharyngeal bulbs was photoconverted. Animals were then rescued from the slide and washed with M9 to dilute the sodium azide. Recovery was assessed using the same imaging settings after allowing the worms to develop normally for 5 hrs. Animals rescued all developed normally and had no morphological defects (n = 20/20 animals).

### Image analysis, processing, and quantification

#### LET-2 and EMB-9 BM fluorescence measurements and protein translation assessment

Single z-slice confocal images were quantified using FIJI (2.0) (Schindelin et al., 2012). BM fluorescence intensity was measured by drawing 3 pixel-wide line along the BM of interest outlined below and recording the mean fluorescence value. Pharyngeal BM measurements were recorded from both sides of the terminal bulb and averaged. Body wall muscle BM measurements were recorded on both sides of the head region flanking the pharynx and averaged. The spermathecal BM was measured on the side proximal to unfertilized eggs. Measurements were recorded on both dorsal and ventral sides of the organ (in regions where it was not folded in on itself) and averaged together. Distal tip cell (DTC) BM measurements were taken around the entire cell. To account for autofluorescence, background measurements were acquired by drawing a line either next to the pharynx (for pharyngeal or body wall muscle BM measurements) or inside the gonad (DTC and spermathecal BM measurements), where no type IV collagen signal was apparent. mNG translational reporter measurements were taken by recording the mean fluorescence value in a single z-slice confocal image for the entire DTC, the entire spermatheca, or two body wall muscles flanking both sides of the pharynx (which were then averaged together). To account for autofluorescence, background measurements were acquired by drawing a line either next to the pharynx (for body wall muscle translation measurements) or inside the vulva (for DTC and spermatheca translation measurements), where no translation reporter signal was apparent.

#### Normalization of data for LET-2/EMB-9 BM protein and translation reporter ratios

In Fig. 4 B, Fig. 6 C, and Fig. 7 C, to directly compare LET-2/EMB-9 ratios between tissues, LET-2::mNG mean fluorescence intensity data points were normalized to the mean value of EMB-9::mNG within that tissue for each trial. Specifically, each data point in the Boxplots was normalized with the following formula: (L / mean E), where “L” is a single mean fluorescence intensity value measurement of LET-2::mNG in a tissue BM and “mean E” is the mean value of all EMB-9::mNG measurements within the same tissue BM for the trial. For Fig. S4 A, Fig. S5 B, and Fig. S5 E the EMB-9::mNG data points were normalized in a similar way to display the spread of data around the mean. Specifically, the formula used was (E / mean E), where “E” is a single measurement of EMB-9::mNG mean fluorescence in a tissue BM. For Fig. 5 B and Fig. S4 D, the same normalization methods were used, substituting measurements of *let-2* translation reporter for “L” and *emb-9* translation reporter for “E”.

#### Normalization of data for RNAi and overexpression experiments

To compare percent loss or gain of EMB-9::mNG or LET-2::mNG in RNAi or overexpression experiments, the data points in Fig. 6 B and Fig. 7 B were normalized by dividing each treatment mean fluorescence value measurement by the mean value of the control measurements within that trial. Specifically, the formula used was (T / mean C) where “T” is a single fluorescence intensity measurement from the treatment animal and “mean C” is the mean value of all measurements from the control animals within a trial. For Fig. S5 A and Fig. S5 D, data points were normalized in a similar way to show the spread of control measurements around the mean. Specifically, the formula used was (C / mean C), where “C” is a single measurement from the control animals.

#### Quantification of type IV collagen muscle aggregates

The number of EMB-9::mScarlet-I (IS1) and EMB-9::mCherry (IS2) fluorescent aggregates that localized within the ER (marked with ER membrane protein ELO-1::mNG) in body wall muscles (Fig. 2 B and Fig. S2 B) was scored in a blinded manner--aggregates were first identified and then it was determined if they were within the ER. A total of 10 aggregates were counted per animal and 10 animals were analyzed for the EMB-9::mScarlet-I (IS1) and EMB-9::mCherry (IS2) strains.

#### FRAP analysis

For FRAP analysis the mean fluorescence intensity values were generated by drawing an ∼0.3μm wide and 3.825μm long line through a clearly defined section of the mid-plane of a single z-slice of the pharyngeal terminal bulb. Background was calculated by drawing an equivalent line within the terminal bulb of the pharynx, where no type IV collagen is produced or localized. Mean fluorescence intensity was measured within the photobleached region and within an equivalent region on the unbleached side of the terminal bulb before bleach (pre-bleach), immediately after bleach (bleach), and 5 h after bleach (5 h post-bleach). A bleach correction factor was used to account for general photobleaching during image acquisition across the duration of the experiment (Gianakas et al., 2023). In brief, the fluorescence intensity measurement in the unbleached region at the 5 h post-bleach timepoint was divided by the value at the pre-bleach timepoint to obtain the bleach correction factor. Fluorescence intensity measurements at the pre-bleach and bleach timepoints in the photobleached region were then multiplied by the bleach correction factor to normalize these values to the 5 h post-bleach timepoint. Background signal was then subtracted from bleach corrected values and used in analysis. Recovery was calculated by dividing the measurement at the 5 h post-bleach timepoint by that at the pre-bleach timepoint.

#### Photoconversion/optical highlighting

For photoconversion analysis, mean fluorescence intensity values were generated by drawing an ∼0.3μm wide and 3.825μm long line through a clearly defined section of the body wall muscle BM located between the two pharyngeal bulbs in a single z-slice image. Background was calculated by drawing an equivalent line next to the pharynx. Mean fluorescence intensity of the red channel was measured within the converted region immediately after photoconversion (conversion) and five hrs after photoconversion (5 h post-conversion). Mean background signal was then subtracted from mean fluorescence intensity values and used for analysis. Percent reduction in photoconverted signal was calculated by dividing the value at the 5 h post-photoconversion timepoint by the value at the conversion timepoint and subtracting that fraction from 1.

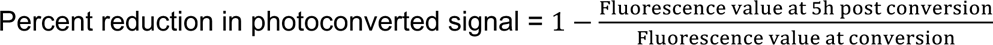

### Time to starvation assay

For each replicate four L4 hermaphrodites were placed on an NGM plate seeded with 400 μL of OP50 E. coli. Plates were maintained at 20°C and checked twice daily for starvation. For every experiment, N2 wild-type animals were plated on the same day, using the same batch of plates. The time it took for experimental strains to starve was reported relative to the time it took the corresponding N2 plates to starve.

### Western blots

Mixed populations of worms were washed off NGM plates with sterile water and placed into 1.5 mL eppendorf tubes. Animals were then washed three times with sterile water and frozen in liquid nitrogen. Frozen animals were then ground to a powder under cryogenic conditions with a mortar and pestle and then resuspended in cooled RIPA buffer plus 1X SIGMAFAST Protease Inhibitor Cocktail (Table S3). Laemmli sample buffer and β-mercaptoethanol (Table S3) were then added to a final concentration of 1X and 4% respectively. The final concentration of sample powder to buffer was 0.5 mg/mL. Samples were boiled for 5 mins and then 15 μL of resuspended samples were loaded onto 4-15% gradient Mini-PROTEAN TGX Gels (Table S3) and run at 100 V. Wet transfers to nitrocellulose were carried out overnight in a cold room at 30 V for 16 hrs. Nitrocellulose membranes were blocked with 5% milk in TBST for 1 h, incubated with primary antibody against mNG in 1% milk and TBST overnight at 4°C (antibody dilution 1:1000), and then treated with secondary peroxidase-conjugated antibody (Table S3) for 1 h at room temperature. Blots were treated with SuperSignal West Atto Ultimate Sensitivity Chemiluminescent Substrate (Table S3) for 5 min before imaging. Blots were imaged on an iBright 1500 imaging system (Invitrogen).

### Sample fractionation and processing for mass spectrometry

Samples were processed by chemical fractionation to enrich for basement membrane (BM) proteins for mass spectrometry-base proteomics as described previously (Morais et al., 2023). Briefly, a mixed population of L1, L2 and L3 larvae were collected in a Tris-lysis buffer (10 mM Tris, 150 mM NaCl, 1% Triton X-100, 25 mM EDTA added of EDTA-free Roche complete protease inhibitor cocktail) and homogenized to extract soluble proteins. After centrifugation at 14,000 × g, the protein supernatant was collected. This fraction is enriched for soluble cellular proteins. The remaining pellet was then resuspended in an alkaline detergent buffer (20 mM NH_4_OH and 0.5% Triton X-100 in 0.1 M PBS) to disrupt cell-matrix bonds, centrifuged again, and the supernatant was removed. The final pellet was resuspended in TEAB/SDS lysis buffer (50 mM TEAB and 5% SDS in LC-MS grade water). This matrix enriched fraction was sonicated with a Covaris LE220+ Focused Ultrasonicator (Covaris). The cellular fraction was added to 2X TEAB/SDS lysis buffer, then both soluble and insoluble protein lysates were reduced with 5 mM dithiothreitol and further alkylated with 15 mM iodoacetamide and processed for overnight digestion at 37°C in S-Trap columns (Protifi) using a Sequencing Grade Modified Trypsin (Promega, Catalog # V5111). The peptide mixtures were eluted from the columns, desalted, and dried to completeness by vacuum centrifugation for mass spectrometry.

### Mass spectrometry data acquisition and analysis

Peptide samples were analyzed by liquid chromatography-tandem mass spectrometry with a Thermo Exploris 480 mass spectrometer (Thermo Fisher Scientific). The raw spectra data were analyzed using MaxQuant version 2.5.0.0 (Cox and Mann, 2008). MS/MS data were searched against a modified version of the Uniprot *Caenorhabditis elegans* proteome (v. 2024_01; containing 26,706 reviewed and unreviewed entries, with the addition of the protein sequence for mRuby2 and mNG (downloaded from https://www.fpbase.org/) using MaxQuant’s integrated search engine, Andromeda. Carbamidomethylation of cysteine was specified as a fixed modification, and oxidation of methionine, proline, and lysine, and *N*-terminal acetylation as variable modifications. Up to two missed tryptic cleavage sites were allowed. Mass tolerance settings for precursor peptide ions were 20 ppm for the first MS/MS search and 4.5 ppm for the main MS/MS search, with 0.5 Da for fragment ions. A target-decoy search strategy was employed to control false-positive identifications, maintaining a 1% false discovery rate (FDR) for both peptide-spectrum matches (PSM) and protein groups. The match-between-runs algorithm was enabled to increase the number of identifications. Contaminants were identified from matches to the MaxQuant contaminants database. Data processing and analysis were performed using R Studio version 2023.9.1.494 (https://rstudio.com). Peptide data obtained with MaxQuant were processed using the QFeatures package version 1.12.0 (Gatto L., 2024). Peptide from decoy and contaminant proteins, non-unique, and detected in only one sample were removed. Peptide intensities were log2-transformed and normalized through median centering. Peptide data were summarized to protein groups via robust summarization, and statistical comparisons were performed with multiple t-tests with Benjamini-Hochberg FDR correction using the msqrob2 package (Goeminne et al., 2016; Goeminne et al., 2020).

### Statistical analysis

Statistical analysis was performed in GraphPad Prism 10. Distribution of data was assessed for normality using Kolmogorov-Smirnov test. To compare fluorescence intensities or peptide levels between two populations, we used an unpaired t-test for normally distributed data or a Mann-Whitney test for data that was not normally distributed. For Fig. 6, we used a t-test with Welch correction because the variance between data sets was unequal. To compare fluorescence intensities between three or more populations, we used a one-way ANOVA followed by Tukey’s multiple comparisons test for normally distributed data with equal variance between groups or a Brown-Forsynth and Welch ANOVA test followed by a Dunnett’s test for normally distributed data with unequal variance between groups. For data that was not normally distributed, we used a Kruskal-Wallis test followed by a Dunn’s test. Sample sizes were determined before each experiment. Sample sizes of at least 10 were chosen for each biological replicate of experiments in Figs 2, 4, 5, 8, Supplementary Figs 2 A, 2 B, and 4 A. Where protein ratios were calculated, three biological replicates of those experiments were carried out to ensure reproducibility of the results. Sex was not a factor in analyzing data because *C. elegans* primarily exist as hermaphrodites. Data points were only excluded if the animal moved during acquisition or developed vacuoles and died.

## Online Supplemental Material

Fig. S1, related to Fig. 1, shows the endogenous locus of fluorophore insertion for all genome edited strains. Flexible linker sequences and Cas9 guide RNA sequences are listed. The fluorophore amino acid (aa) length and aa sequence similarity between fluorophores are shown. Fig. S2, related to Fig. 2, shows intracellular aggregates of LET-2::mNG (C) and EMB-9::mNG (C) in the body wall muscles of *C. elegans* as they develop from L1 larvae to gravid adults. Also shown are the localization of EMB-9::mCherry (IS2) aggregates within the ER and FRAP analysis of LET-2::mNG (C), EMB-9::mNG (C), and EMB-9::mNG (IS1) at the pharyngeal BM. Fig. S3, related to Fig. 3, shows the abundance of mNG and mRuby2 proteins determined by LC MS/MS in the larval cellular fractions of strains expressing the knock-in proteins Strain 1 [EMB-9::mRuby2 (C); LET-2::mNG (C)] and Strain 2 [EMB-9-mNG (C); LET-2::mRuby2 (C)]. Also shown are the comparable abundances of LET-2 and EMB-9 protein in the matrix fractions of both strains. Fig. S4, related to Fig. 4 and 5, shows the normalized LET-2::mNG (C) data with the normalized EMB-9::mNG (C) data and the normalized LET-2 translation reporter (*let-2::P2A::PEST::mNG*) data with the normalized EMB-9 translation reporter (*emb-9::P2A::PEST::mNG*) data for all tissues and trials. Also shown is a western blot confirming the stability of mNG associated with the LET-2::mNG (C) and EMB-9::mNG (C) fusion proteins and mean fluorescence intensity measurements of both LET-2::mNG (C) and EMB-9::mNG (C) at the pharyngeal BM of homozygotes and heterozygotes. Fig. S4, related to Figs. 6, 7, and 8, shows the normalized LET-2::mNG (C) data with the normalized EMB-9::mNG (C) data and the normalized treatment (RNAi or overexpression) data with the normalized control (L4440 empty vector RNAi or absence of overexpression) data for all trials. Also shown are intracellular aggregates of LET-2::mNG (C) and EMB-9::mNG (C) in the muscles of day 1 adult *C. elegans* after *emb-9* RNAi and *emb-9* overexpression. Additionally, sum projections from confocal z-stacks of L1 larvae carrying *let-2* or *emb-9* glycine mutant alleles grown at restrictive temperature are shown. Table S1 outlines the strains used in this study. Table S2 shows the primer sequences used in this study. Table S3 lists the chemical and DNA reagents used.

## Data Availability

Plasmids generated, *C. elegans* strains generated, and data are available upon request. Strains will be donated Caenorhabditis Genetics Center (CGC). The mass spectrometry proteomics data have been deposited to the ProteomeXchange Consortium via the PRIDE partner repository (Perez-Riverol et al., 2022) with the dataset identifier PXD055575 and 10.6019/PXD055575. All other data are included in the manuscript and supplemental materials.

## Acknowledgements

We thank S. Boudko for extensive discussions on type IV collagen biochemistry, R. Jayadev for valuable suggestions when conceptualizing the project, and members of the Sherwood lab for feedback on experimental analysis. We also thank G. Poulin for help maintaining and handling *C. elegans* strains used for mass spectrometry. Some strains were provided by the CGC funded by NIH Office of Research Infrastructure Programs (P40 OD010440). This work was supported by NIH grants (R35GM118049, R21OD032430) and a Wellcome Trust Discovery Award (Ref: 227417/Z/23/Z) to D.R. Sherwood, the NIH Ruth L. Kirschstein Individual Postdoctoral Fellowship (F32AG077907) to W. Ramos-Lewis and the Jane Coffin Childs Memorial Fund for Medical Research to A.W.J. Soh. R. Lennon was supported by the Wellcome Trust (Ref: 226804/Z/22/Z, Ref: 227417/Z/23/Z, Ref: 301803/Z/23/Z), Kidney Research UK and The Stoneygate Trust (Alport Research Hub) and the National Institute for Health Research (NIHR-Manchester BRC).

The authors declare no competing financial interests.

## Author Contributions

S. Srinivasan, W. Ramos-Lewis, and D.R. Sherwood conceived the project. S. Srinivasan and W. Ramos-Lewis built strains and molecular constructs, conducted experiments, analyzed the data, and interpreted the data, with the following contributions from other authors: M.R.P.T. Morais, E. Williams and R. Lennon designed and reported the proteomics experiments presented in Fig. 3A and Fig. S3. E. Williams prepared the proteomics samples for analysis and M.R.P.T. Morais performed data analysis and prepared the figures. Q. Chi designed and built *C. elegans* strains used in Figs. 2, 3, 4, 6, 7 and 8, and Figs. S2, S3, S4 and S5. A.W.J. Soh helped design data analysis techniques. S. Srinivasan and W. Ramos-Lewis prepared all other figures. D.R. Sherwood, S. Srinivasan, and W. Ramos-Lewis prepared the manuscript. D.R. Sherwood, S. Srinivasan, W. Ramos-Lewis, R. Lennon, M.R.P.T. Morais, and E. Williams edited the manuscript.

## Supplemental figure legends

Figure S1. **Knock-in sites and comparisons between different genetically encoded fluorophores (A)** The schematic illustrates the *C. elegans* type IV collagen genes *emb-9* and *let-2*. Exons are depicted as blue boxes. The lines indicate the endogenous sites where the indicated fluorophores were inserted, and listed underneath are the two amino acids the fluorophore was inserted between. The precise nucleic acid sequence at the insertion site and translated amino acid sequence are listed. Linkers used to flank fluorophores are depicted as wavy black lines and their sequences are listed. For each knock-in site, the sgRNA sequence(s) and PAM site used to target the gene for CRISPR mediated homologous recombination are shown. Strains with mNG knocked-in at the *let-2* N-terminal site (N) or *let-2* Internal Site 1 (IS1) were homozygous inviable and were not further evaluated in this study. **(B)** Left: The table shows the amino acid (aa) length of each fluorophore used in this study. Right: The table shows the aa sequence similarity between each pair of fluorophores.

Figure S2. **Internalization and FRAP analysis of LET-2 and EMB-9 knock-ins (A)** Representative sum projections of confocal fluorescence z-stacks of LET-2::mNG (C) and EMB-9::mNG (C) at the L1, L2, L3, Late L4 and gravid adult stages. Small aggregates in the body wall muscle were first detected in L3 larvae (47% of LET-2::mNG (C) and 73% of EMB-9::mNG (C), n ≥ 10 each) and continued into gravid adult (100% for LET-2::mNG and EMB-9::mNG, n ≥ 10 each). The scale bar is 20 μm. **(B)** Single z-slice fluorescence image of EMB-9::mCherry (IS2) muscle aggregates (magenta in merge) localized within the ER (visualized with ELO-1::mNG; cyan in merge) in L1 larvae. Yellow arrows point to aggregates in the ER (n = 10 animals; 10 aggregates counted per animal; 99/100 aggregates localized to the ER). The scale bar is 5 μm. **(C)** Left: Single z-slice images of LET-2::mNG (C), EMB-9::mNG (C), and EMB-9::mNG (IS1) prior to, immediately after, and 5h after FRAP at the L1 proximal pharyngeal bulb. Yellow arrowheads point to photobleached regions. White arrows point to unbleached regions used as the control region for analysis. Dotted boxes indicate the location of magnified regions shown in the panel below. The scale bar is 1μm for magnified panels. Scale bar is 10 μm for all other panels. Right: Boxplots show the percent of fluorescent signal recovered after 5 h (n <u>=</u> 10 animals, p > 0.05, ANOVA with Tukey’s multiple comparisons).

Figure S3. **LET-2 is approximately twice as abundant as EMB-9 in larva and the total EMB-9 and LET-2 protein is comparable between strains (A,B)** Boxplots display the abundance of mNG and mRuby2 represented by their respective log2-protein intensity (a.u.) determined using LC-MS/MS. The samples displayed are the larval (L1-L3 stages) cellular fractions of Strain 1 [EMB-9::mRuby2 (C); LET-2::mNG (C)] and Strain 2 [EMB-9-mNG (C); LET-2::mRuby2 (C)]. The log2-fold changes from the comparison between strains are listed above the boxplots. The mNG (LET-2) in Strain 1 (n=3) was approximately 1.7 times the abundance of mNG (EMB-9) in Strain 2 (n=4) while mRuby2 (LET-2) was approximately 1.7 times more abundant in Strain 2 (n=3) than mRuby2 (EMB-9) in Strain 1 (n=2). **(C,D)** Boxplots display the abundance of EMB-9 and LET-2 represented by their respective log2-protein intensity (a.u.). The samples displayed are the larval matrix fractions of Strain 1 and Strain 2 (n=5 each). log2-fold changes from the comparison between strains are listed as well.

Figure S4. **Normalization of LET-2/EMB-9 ratios, absence of quenching, and stability of fluorophore tagging at the C-terminus (A)** The boxplots show the normalized LET-2::mNG (C) data with the normalized EMB-9::mNG (C) data for every tissue shown in Fig. 4 B. Colors of the data points indicate different trials. Normalization was performed by dividing the mean BM fluorescence intensity measurement of each LET-2::mNG (C) or EMB-9::mNG (C) animal by the average of all mean BM fluorescence intensity measurements of EMB-9::mNG (C) for the same tissue and trial (see Methods, n ≥ 10 animals for each of three trials). **(B)** Western blot using anti-mNG primary antibody on non-sonicated total lysate from *let-2::mNG (C)* and *emb-9::mNG (C)* animals (LET-2::mNG and EMB-9::mNG protein), transgenic animals expressing mNG driven by promoter for *let-2* (*let-2p>mNG*), and wild-type N2 animals lacking mNG. Red arrows indicate predominant bands in each lane. **(C)** Top: Single z-slice confocal fluorescence images of pharynxes from late L4 staged EMB-9::mNG (C) and LET-2::mNG (C) homozygotes and heterozygotes. The scale bar is 20 μm. Bottom: The boxplots show the mean fluorescence intensity measurements from the BM surrounding the proximal pharyngeal bulb (n≥10 animals). **(D)** The boxplots show the normalized LET-2 translation reporter (*let-2::P2A::PEST::mNG*) data with the normalized EMB-9 translation reporter (*emb-9::P2A::PEST::mNG*) data plotted for each tissue shown in Fig. 5 B. Colors of the data points indicate different trials. Normalization was performed by dividing the mean BM fluorescence intensity measurement of each LET-2 translation reporter or EMB-9 translation reporter animal by the average of all mean BM fluorescence intensity measurements of EMB-9 translation reporter for the same tissue and trial (see Methods, n ≥ 10 animals for each of three trials).

Figure S5. **Normalization of LET-2/EMB-9 ratios and internal aggregates after *emb-9* reduction or overexpression and in *let-2* or *emb-9* glycine substitution mutants (A)** The boxplots show the normalized *emb-9* RNAi knockdown and control RNAi (L4440 empty vector) data in LET-2::mNG (C) and EMB-9::mNG (C) animals shown in Fig. 6 B. Colors of the data points indicate different trials. Normalization was performed by dividing the mean BM fluorescence intensity (LET-2::mNG (C) or EMB-9::mNG (C)) measurement from each *emb-9* RNAi or control animal by the average of all mean BM fluorescence intensity measurements from control animals for the same trial (see Methods, n ≥ 8 animals for each of three trials, t-tests with Welch correction, **** p-value < 0.0001). **(B)** The boxplots show the normalized LET-2::mNG (C) and normalized EMB-9::mNG (C) data for control RNAi (L4440 empty vector) and *emb-9* RNAi knockdown animals shown in Fig. 6 C. Colors of the data points indicate different trials. Normalization was performed by dividing the mean BM fluorescence intensity measurement of each LET-2::mNG (C) or EMB-9::mNG (C) animal by the average of all mean BM fluorescence intensity measurements of EMB-9::mNG (C) for the same trial (see Methods, n ≥ 8 animals for each of three trials). **(C)** Single z-slice images of LET-2::mNG (C) (left) and EMB-9::mNG (C) (right) in day 1 adult worms (72h post hatch) with and without *emb-9* RNAi knockdown. Yellow arrowheads indicate body wall muscle intracellular aggregates. Increased intracellular aggregation was seen in LET-2::mNG (C) *emb-9* RNAi treated worms and decreased intracellular aggregation was seen in EMB-9::mNG (C) *emb-9* RNAi treated worms (n = 28/28 worms observed for each). Scale bar is 10μm. **(D)** The boxplots show the normalized *emb-9* overexpression and control (no overexpression) data in LET-2::mNG (C) and EMB-9::mNG (C) animals shown in Fig. 7 B. Colors of the data points indicate different trials. Normalization was performed by dividing the mean BM fluorescence intensity (LET-2::mNG (C) or EMB-9::mNG (C)) measurement of each overexpression or control animal by the average of all mean BM fluorescence intensity measurements of control animals for the same trial (see Methods, n ≥ 5 animals for each of three trials, unpaired t-test (left) and a t-test with Welch correction (right), **** p-value < 0.0001, ns, not significant). **(E)** The boxplots show the normalized LET-2::mNG (C) and normalized EMB-9::mNG (C) data after *emb-9* overexpression and in control (no overexpression) animals shown in Fig. 7 C. Colors of the data points indicate different trials. Normalization was performed by dividing the mean BM fluorescence intensity measurement of each LET-2::mNG (C) or EMB-9::mNG (C) animal by the average of all mean BM fluorescence intensity measurements of EMB-9::mNG (C) for the same trial (see Methods, n ≥ 5 animals for each of three trials). **(F)** Single z-slice images of LET-2::mNG (C) (left) with and without *emb-9::mRuby2* overexpression and EMB-9::mNG (C) (right) with and without *emb-9::mNG::mRuby* overexpression in day 1 adult worms (72h post hatch). Yellow arrowheads indicate body wall muscle intracellular aggregates. Increased intracellular aggregation was seen in EMB-9::mNG (C) worms with *emb-9::mNG::mRuby* overexpression in all body wall muscle segments (n=11/11 animals observed). Scale bar is 10μm. **(G)** Representative sum projections of confocal z-stacks of LET-2::mNG (C) and EMB-9::mNG (C) wild-type animals, LET-2::mNG G1287D and EMB-9::mNG (C) G1173D mutant animals, and animals with a wild-type fluorescently tagged protein paired with an untagged mutant protein: EMB-9 G1173D; LET-2::mNG (C) and EMB-9::mNG (C); LET-2 G1173D. The imaged L1 larvae were grown at 25°C (restrictive temperature for *emb-9* and *let-2* mutants). Increased intracellular aggregation was seen in animals carrying *let-2* or *emb-9* glycine mutant alleles (n=20/20 animals per condition). Scale bar is 10μm.

Table S1. ***C. elegans* strains used in this study**

Table S2. **Oligonucleotide sequences used in this study**

Table S3. **Chemicals and DNA reagents used in this study**

