## Supplemental Figures and Tables for "A complete collagen IV fluorophore knock-in toolkit reveals α-chain diversity in basement membrane"

Supplementary Figure 1

A

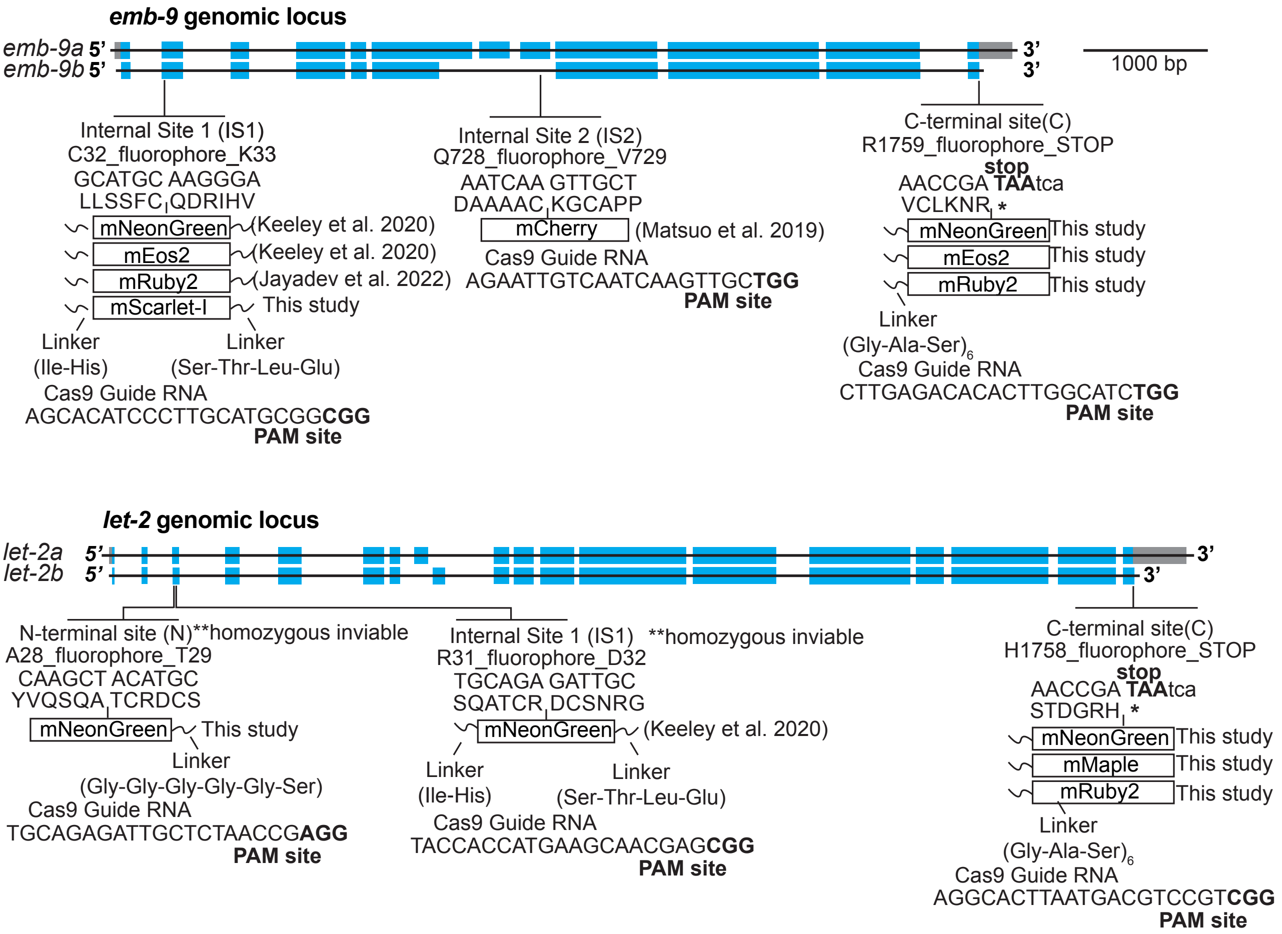


B

| Fluorophore | Length (aa) |
|---|---|
| mNG | 236 |
| mEos2 | 226 |
| mMaple | 237 |
| mScarlet-I | 232 |
| mCherry | 236 |
| mRuby2 | 237 |

| Fluorophore | mNG | mEos2 | mMaple | mScarlet-I | mCherry | mRuby2 |
|---|---|---|---|---|---|---|
| mNG | 100.00 | 24.18 | 29.92 | 29.10 | 31.15 | 27.05 |
| mEos2 | | 100.00 | 74.59 | 48.36 | 44.67 | 48.36 |
| mMaple | | | 100.00 | 54.92 | 53.69 | 50.82 |
| mScarlet-I | | | | 100.00 | 85.66 | 59.84 |
| mCherry | | | | | 100.00 | 55.74 |
| mRuby2 | | | | | | 100.00 |

Supplementary Figure 2

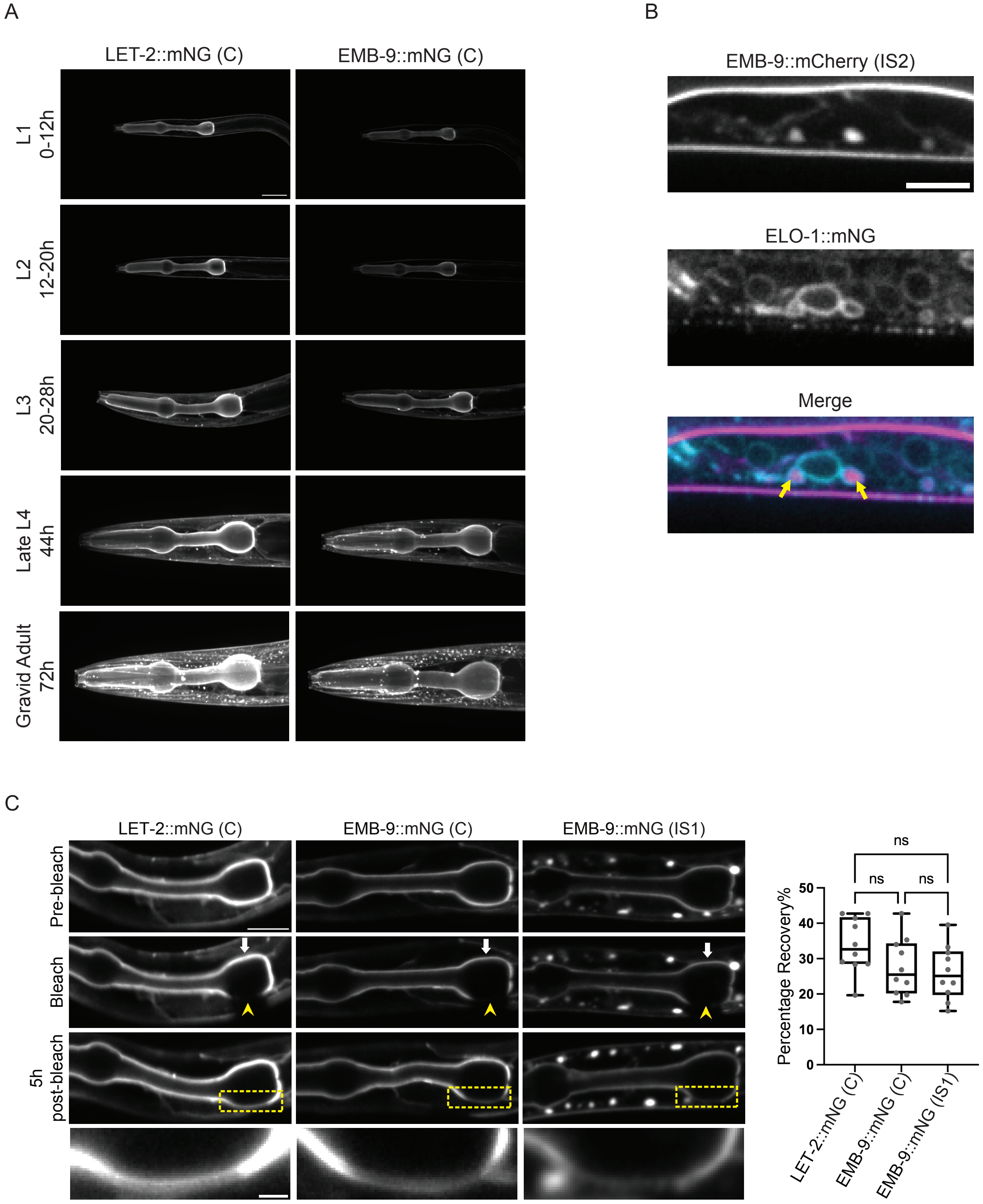

Supplementary Figure 3

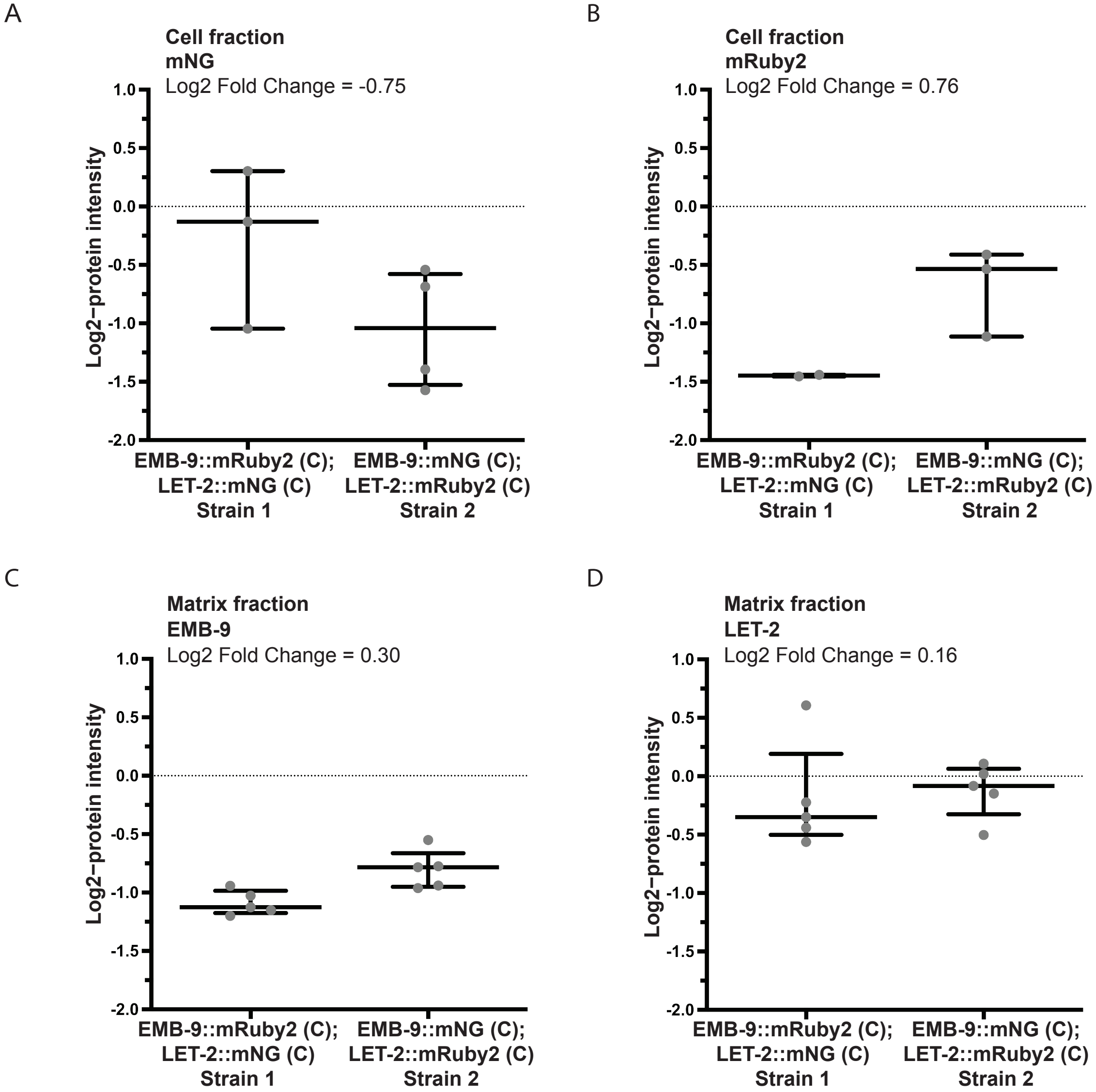

Supplementary Figure 4

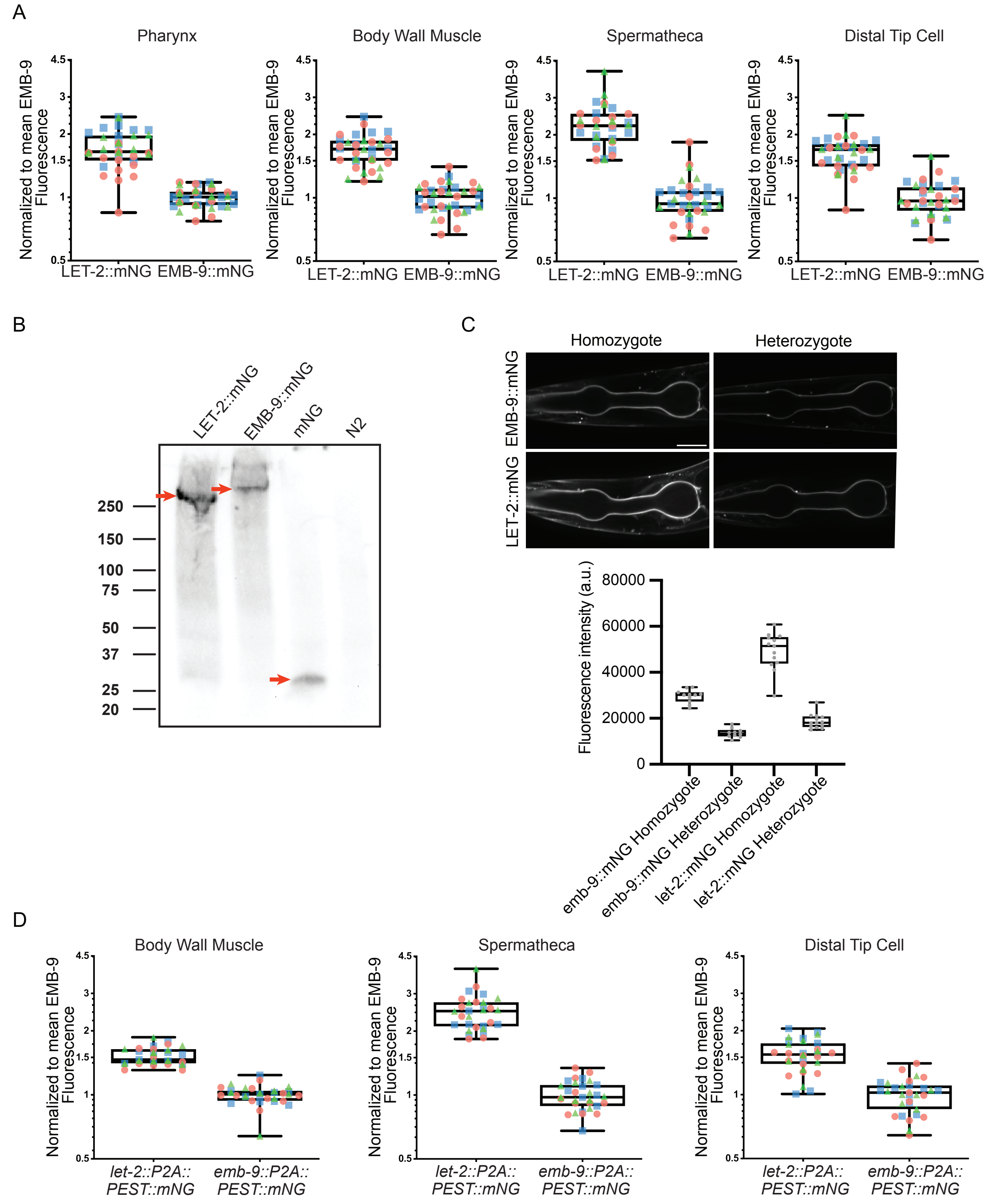

Supplementary Figure 5

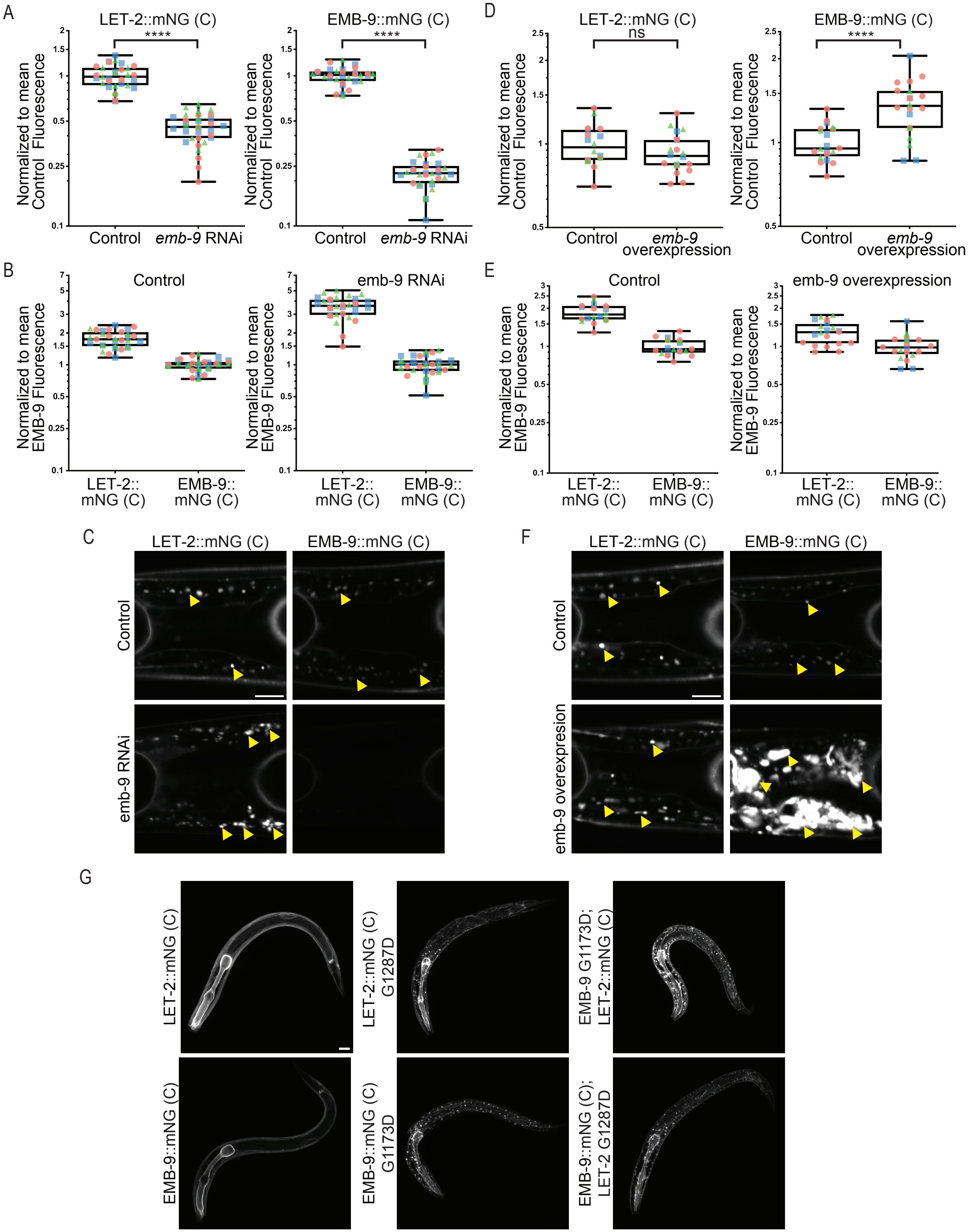

| Table S1. ***C. elegans* strains used in this study** | | | |
|---|---|---|---|
| Strain | Genotype | Fluorophore Location * | Reference |
| N2 | Wild-type (ancestral) | | |
| IHR169 | *xyz9 [emb-9::mCherry]* III | IS2 | Matsuo et al., 2019 |
| NK2326 | *qy24 [emb-9::mNG]* III | IS1 | Keeley et al., 2020 |
| NK2604 | *qy89 [emb-9::mEos2]* III | IS1 | Keeley et al., 2020 |
| NK2585 | *qy83 [emb-9::mRuby2]* III | IS1 | Jayadev et al., 2022 |
| NK2762 | *qy152 [emb-9::mScarlet-I]* III | IS1 | This study |
| NK3057 | *qy236 [emb-9::mNG]* III | C | This study |
| NK3026 | *qy228 [let-2::mNG]* X | C | This study |
| NK3072 | *qy244 [emb-9::mRuby2]* III | C | This study |
| NK2984 | *qy216 [let-2::mRuby2]* X | C | This study |
| NK3077 | *qy244 [emb-9::mRuby2]* III; *qy228 [let-2::mNG]* X | C | This study |
| NK3083 | *qy236 [emb-9::mNG]* III; *qy216 [let-2::mRuby2]* X | C | This study |
| NK3073 | *qy245 [emb-9::mEos2]* III | C | This study |
| NK3242 | *qy290 [let-2::mMaple]* X | C | This study |
| NK3248 | *qy152 [emb-9::mScarlet-I]* III*; qy97 [elo-1::mNG]* IV | C | This study |
| NK3249 | *xyz9 [emb-9::mCherry]* III*; qy97 [elo-1::mNG]* IV | IS2 | This study |
| NK3236 | *qy252 [let-2p>mNG]* I | n/a | This study |
| NK3237 | *qy286 [let-2::P2A::PEST::mNG]* X | n/a | This study |
| NK3238 | *qy287 [emb-9::P2A::PEST::mNG]* III | n/a | This study |
| NK3094 | *b117 [emb-9(G1173D); qy228 [let-2::mNG]* | C | This study |
| NK3095 | *qy236 [emb-9::nNG]* III*; b246 [let-2(G1287D)]* X | C | This study |
| NK3240 | *qy288 [emb-9 (G1173D)::mNG]* III | C | This study |
| NK3241 | *qy289 [let-2 (G1287D)::mNG]* X | C | This study |
| NK3239 | *qy245 [emb-9::mEos2]* III*; b246 [let-2(G1287D)]* X | C | This study |
| NK3243 | *b117 [emb-9(G1173D)]* III*; qy290 [let-2::maple]* X | C | This study |

* Refers to the position of fluorescent protein within the coding region of type IV collagen genes
n/a = not applicable

Table S2. **Oligonucleotide sequences used in this study**

| Identifier | | | | Oligonucleotide sequence (5' → 3') | Template |
|---|---|---|---|---|---|
| Gene | Insertion | Site | Type | | |
| Genome-edited Strains Cas9 Short Guide Primers | | | | | |
| *let-2* | mNG, mRuby2, mMaple | C | Forward | cctcctattgcgagatgtcttGaggcacTTAATGACGTCCGTGTTTTAGAG CTAGAAATAGCA | pDD122 (Dickinson et al., 2013) |
| *emb-9* | mNG, mRuby2, mEos2 | C | Forward | cctcctattgcgagatgtcttGCTTGAGACACACTTGGCATCGTTTTAG AGCTAGAAATAGCAAG | pDD122 (Dickinson et al., 2013) |
| *emb-9* | mScarlet-I | IS1 | Forward | tcctattgcgagatgtcttGAGCACATCCCTTGCATGCGGGTTTTAGA GCTAGAAATAGC | pDD122 (Dickinson et al., 2013) |
| Genome-edited Strains Genotyping Primers | | | | | |
| *let-2* | mNG, mRuby2, mMaple | C | Forward | CTCCAAAGATTCTCCACCAT | *genomic* DNA |
| *let-2* | mNG, mRuby2, mMaple | C | Reverse | CCTTGTCCGACCATGTCGAA | *genomic* DNA |
| *emb-9* | mNG, mRuby2, mEos2 | C | Forward | AGTACCACAGTGCCCACCAGGA | *genomic* DNA |
| *emb-9* | mNG, mRuby2, mEos2 | C | Reverse | CCTTGTCCGACCATGTCGAA | *genomic* DNA |
| *emb-9* | mScarlet-I | IS1 | Forward | TGATAAATAGGTGGAGCGCGAT | *genomic* DNA |
| *emb-9* | mScarlet-I | IS1 | Reverse | TCCTGGATCTCCCTTTGCTC | *genomic* DNA |
| Genome-edited Strains Repair template Primers | | | | | |
| *let-2** | mNG, mRuby2, mMaple | C | Forward | gtaatacgactcactatagggcgaattgggtaccacaactagtGAGAGAAGGGAAT GGGAGGT | *genomic* DNA |
| *let-2** | mNG, mRuby2, mMaple | C | Reverse | CTCCCGATGCTCCtgAgGCaCCcGAtGCaCCtGAgGCaCCATG ACGTCCGTCGGTGGACTTGA | *genomic* DNA |
| *let-2** | mNG, mRuby2, mMaple | C | Forward | AGCATACATTATACGAAGTTATTTTCAGGGAGCCGGATCTT AAGTGCCTACTCACTTTACTTAG | *genomic* DNA |
| *let-2** | mNG, mRuby2, mMaple | C | Reverse | AAGGGAACAAAAGCTGGAGCTCCAGCGGCCGCTTTGCATG CCACTACTAGCAACTGAATTATAG | *genomic* DNA |
| *emb-9** | mNG, mRuby2 | C | Forward | CGCGTAATACGACTCACTATAGGGCGAATTGGGTACCACA ACTAGTCAGTGGACAAGATCTTGGTCAA | *genomic* DNA |
| *emb-9** | mNG, mRuby2 | C | Reverse | TGCTCCtgAgGCaCCcGAtGCaCCtGAgGCaCCTCGaTTCTTG AGACACACTTGGCATCTtGATACACG | *genomic* DNA |
| *emb-9** | mNG, mRuby2 | C | Forward | GCATACATTATACGAAGTTATTTTCAGGGAGCCGGATCTTA ATCATCCCAATCATCACTCGC | *genomic* DNA |
| *emb-9** | mNG, mRuby2 | C | Reverse | GGGAACAAAAGCTGGAGCTCCAGCGGCCGCTTTGCATGC TACTTTGAACGTATAACCTC | *genomic* DNA |
| *emb-9* | mEos2 | C | Forward | CGCGTAATACGACTCACTATAGGGCGAATTGGGTACCACA ACTAGTCAGTGGACAAGATCTTGGTCAA | *genomic* DNA |
| *emb-9* | mEos2 | C | Reverse | GGCACCGGAGGCACCGGAGGCTCCGGAGGCTCCTCGaTT CTTGAGACACACTTGGCATCTtGATACACG | *genomic* DNA |
| *emb-9* | mEos2 | C | Forward | GCATACATTATACGAAGTTATTTTCAGGGAGCCGGATCTTA ATCATCCCAATCATCACTCGC | *genomic* DNA |
| *emb-9* | mEos2 | C | Reverse | GGGAACAAAAGCTGGAGCTCCAGCGGCCGCTTTGCATGC TACTTTGAACGTATAACCTC | *genomic* DNA |
| *emb-9* | mScarlet-I | IS1 | Forward | ATACGACTCACTATAGGGCGAATTGGGTACCACAACTAGTT GAATGTCAATTCTCCAGAA | *genomic* DNA |
| *emb-9* | mScarlet-I | IS1 | Reverse | GGCCTCTCCCTTGGAGACCATGTGGATaCAcGCtGCaGCtG CgT | *genomic* DNA |
| *emb-9* | mScarlet-I | IS1 | Forward | GCATACATTATACGAAGTTATTTTCAGGGAGCCGGATCTTC GACACTCGAGAAaGGtTGc | *genomic* DNA |
| *emb-9* | mScarlet-I | IS1 | Reverse | AGGGAACAAAAGCTGGAGCTCCAGCGGCCGCTTTGCATG CGCGTCCTTGTGAATTGACAC | *genomic* DNA |
| Primers used to build the translational reporter constuct | | | | | |
| n/a | P2A build | n/a | Forward | CCGGCCTGCTTCAGCAGGGAGAAGTTGGTGGCGCCTCCG GCTCCCTTGTAGAGCTCGTCCATTC | n/a |
| n/a | P2A build | n/a | Reverse | TTAATGAGCTCGGAGACCATGGGGCCGGGGTTCTCCTCC ACGTCGCCGGCCTGCTTCAGCAGGGAG | n/a |
| n/a | P2A amplify | n/a | Forward | CGGGTGCCTCAGGAGCATCGGGAGCCTCAGGAGCATCGG GCGCCACCAACT | n/a |
| n/a | P2A amplify | n/a | Reverse | GGGGCCGGGGTTCTCCTCCAC | n/a |

| | | | | | |
|---|---|---|---|---|---|
| n/a | PEST | n/a | Forward | CAGGCCGGCGACGTGGAGGAGAACCCCGGCCCCGGAGCCGGACTTAGCCATGGCTTCCCGCCGG | pCFJ150 (Kaymak et al., 2016) |
| n/a | PEST | n/a | Reverse | CTGGGAGGGAGGCCATGTTGTCCTCCTCTCCCTTGGAGACCATCACATTGATCCTAGCAGAAGC | pCFJ150 (Kaymak et al., 2016) |
| Transgenic Construct Primers | | | | | |
| *let-2* | promoter | n/a | Forward | CGACGTTGGCGTCGATCATCCTGTAAAACGACGGCCAGTGCTAGCAGTTGTGCACTTTGTGCAAC | *genomic* DNA |
| *let-2* | promoter | n/a | Reverse | GAGGCCATGTTGTCCTCCTCTCCCTTGGAGACCATTTTGGGCCCGGTGGTAGGCAGCCGCTGGC | *genomic* DNA |
| *emb-9* | *emb-9p::emb-9:: mRuby2 ::emb-9 3'utr* | n/a | Forward | ACGTTGGCGTCGATCATCCTGTAAAACGACGGCCAGTGCGGCCGCAACAACAATCGATTCGGATCAT | *genomic* DNA |
| *emb-9* | *emb-9p::emb-9:: mRuby2 ::emb-9 3'utr* | n/a | Reverse | TCAAGCCGAGAAGCGATAAGCGTGACATCTTCCAGTCACGGCGATCGACT | *genomic* DNA |
| *emb-9* | *emb-9p::emb-9:: mRuby2 ::emb-9 3'utr* | n/a | Forward | AGTCGATCGCCGTGACTGGAAGATGTCACGCTTATCGCTTCTCGGCTTGA | *genomic* DNA |
| *emb-9* | *emb-9p::emb-9:: mRuby2 ::emb-9 3'utr* | n/a | Reverse | TCAACACCTGGTTGTCCAGCAACTTGATTGACAATTCTCAATGGTG | *genomic* DNA |
| *emb-9* | *emb-9p::emb-9:: mRuby2 ::emb-9 3'utr* | n/a | Forward | CACCATTGAGAATTGTCAATCAAGTTGCTGGACAACCAGGTGTTGA | *genomic* DNA |
| *emb-9* | *emb-9p::emb-9:: mRuby2 ::emb-9 3'utr* | n/a | Reverse | TGATTGGGATGATTATCGGTTCTTGAGACACACTTGGCATCTG | *genomic* DNA |
| *emb-9* | *emb-9p::emb-9:: mRuby2 ::emb-9 3'utr* | n/a | Forward | CAGATGCCAAGTGTGTCTCAAGAACCGATAATCATCCCAATCA | *genomic* DNA |
| *emb-9* | *emb-9p::emb-9:: mRuby2 ::emb-9 3'utr* | n/a | Reverse | CAGATGCCAAGTGTGTCTCAAGAACCGATAATCATCCCAATCA | *genomic* DNA |

> Lowercase letters indicate mutations made to repair template to prevent Cas9 cutting.

* knock-in glycine mutation alleles were generated using the respective mNG primers.

n/a Not applicable

Table S3. **Chemicals and DNA reagents used in this study**

| Reagent | Source / Reference | Identifier |
|---|---|---|
| Ampicillin | Sigma-Aldrich | Catalog #A0166 |
| IsopropylIsopropyl β-D-1-thiogalactopyranoside (IPTG) | Sigma-Aldrich | Catalog #I6758 |
| Hygromycin B | Sigma-Aldrich | Catalog #H3274 |
| Sodium azide | Sigma-Aldrich | Catalog #S2002 |
| EcoRI | NEB | Catalog #R0101S |
| ClaI | NEB | Catalog #R0197S |
| RIPA Buffer | Sigma-Aldrich | Catalog #R0278 |
| 1X SIGMAFAST Protease Inhibitor Cocktail | Sigma-Aldrich | Catalog #S8830 |
| Laemmli sample buffer | Bio-Rad Laboratories | Catalog #161-0737 |
| β-mercaptoethanol | Thermo Fisher Scientific | Catalog #21985023 |
| 4-15% gradient Mini-PROTEAN TGX Gels | Bio-Rad Laboratories | Catalog #4561084 |
| SuperSignal West Atto Ultimate Sensitivity Chemiluminescent Substr | Thermo Fisher Scientific | Catalog #A38555 |
| TRIS-buffered saline (TBS-20X) with 2% Tween 20 (TBST) | Thermo Fisher Scientific | Catalog #J60497.K3 |
| **Antibodies** | | |
| 32F6 mouse anti-mNG antibody | ChromoTek | RRID: AB_2827566 |
| Goat anti-Mouse IgG (H+L) Secondary Antibody, HRP | Invitrogen | Catalog #62-6520 |
| **Plasmids** | | |
| pDD122 | Dickinson et al., 2013 | RRID: Addgene_47550 |
| pCFJ352 | Addgene | RRID: Addgene_30539 |
| pAP088 | Pani and Goldstein, 2018 | |
| pCFJ150 | Addgene | RRID: Addgene_19329 |
| pCFJ90 | Addgene | RRID: Addgene_19327 |
| pCFJ104 | Addgene | RRID: Addgene_19328 |
